# Floral humidity as a signal – not a cue – in a nocturnal pollination system

**DOI:** 10.1101/2022.04.27.489805

**Authors:** Ajinkya Dahake, Piyush Jain, Caleb Vogt, William Kandalaft, Abraham Stroock, Robert A. Raguso

**Affiliations:** Department of Neurobiology and Behavior, Cornell University, Ithaca, NY 14853 USA; Sibley School of Mechanical and Aerospace Engineering, Cornell University, Ithaca, NY 14853 USA; Smith School of Chemical and Biomolecular Engineering, Cornell University, Ithaca, NY 14853 USA

## Abstract

Although visual and olfactory floral signals attract pollinators from a distance, at the flower’s threshold, pollinators can use floral humidity as an index cue for nectar presence. We evaluate the role of floral humidity in the *Datura wrightii-Manduca sexta* nocturnal pollination system. In addition to our finding that *M. sexta* shows strong innate attraction toward humid flowers, we identify the hygrosensing sensillum on their antennae, demonstrate its extreme sensitivity to minute changes in RH, and observe the elimination of moths’ behavioral preference towards humid flowers following experimental occlusion of the sensilla. Despite Manduca’s attraction toward humid flowers, we find that floral humidity is not a reliable cue for nectar presence in this system. While Datura floral headspace sustains an enormous humidity gradient, it is not a consequence of nectar evaporation, but an outcome of gas exchange through floral stomata and is decoupled from nectar presence. Using interdisciplinary tools, we demonstrate the function of floral humidity as an attractive signal, not a cue, in this pollination system, thus showcasing an underappreciated modality by which flowers may manipulate their visitors.

## Introduction

The recent expansion of studies in plant-pollinator interactions reflects increased recognition of their importance to ecosystem services, agricultural food security and biological diversity ^1-3^. Examining plant-pollinator interactions through a neuroethological lens presents unique opportunities to investigate the sender-receiver elements of ecological communication ^4,5^, the findings of which have informed current agricultural practices ^6^, and improved integrated-pest-management ^7^.

Plant-pollinator interactions are maintained through floral signals and the innate responses they evoke in the target pollinator. Floral signals are often complex, yet most research has focused on the role of floral secondary metabolites such as volatiles and color pigments in attracting pollinators (but see exceptions: acoustic reflection^8^, nectar guides ^9-11^, heat ^12^). However, the spatial scale at which pollinators are attracted to floral signals and other associated cues is an ongoing topic of investigation. Although floral scent and color can attract pollinators from meters distance ^13-17^, they cease to be informative once pollinators are at a flower’s threshold (mm to cm distance). Recently emptied flowers often remain scented, turgid, and pigmented minutes to hours after nectar was extracted by an earlier visitor. Therefore, to minimize the cost of foraging on empty flowers, pollinators might benefit from more reliable information as they navigate a patch of flowering plants ^18,19^. Floral primary metabolism and transpiration produce gradients in the concentrations of carbon dioxide (CO_2_) and humidity (RH) within the headspace of the flower (mm to cm distance) and may offer more reliable information on nectar availability before pollinators commit to probing or visiting a flower ^20,21^.

Current evidence on floral CO_2_ and RH points to similar roles as index cues for nectar presence and profitability assessment by pollinators. However, the distinction between signals and cues may blur when multiple lines of evidence show that the interests of senders and receivers align, as posited in the sensory drive theory ^22,23^. In plant-pollinator interactions, floral traits are often described interchangeably as signals and cues which may oversimplify their role in this complex inter-kingdom communication ^24^. Signals must be beneficial (on average) for both senders and receivers, improve their fitness, and incur a signaling cost ^25,26^. Cues on the other hand often benefit only one party and may not incur a signaling cost.

In the context of conventional (non-deceptive) pollination, floral CO_2_ decays to ambient levels within hours after anthesis and may serve early visitors as a cue for newly opened flowers ^27^. Flowers with above ambient CO_2_ are more attractive to pollinators than flowers with ambient CO_2_ ^28,29^. However, in the absence of differential profitability between the two flowers, pollinator preference decays to chance over subsequent visits ^30^. Thus, flower-visiting insects can utilize above-ambient floral CO_2_ as an ephemeral profitability cue for freshly opened flowers, due to correlation with unexploited floral rewards.

Unlike CO_2_, floral humidity indicates nectar presence to foraging pollinators as a direct physical consequence of nectar evaporation, rather than as a correlated aspect of anthesis ^31^; it may therefore alert pollinators to flowers that refill nectar after anthesis. The evening primrose flower (*Oenothera cespitosa)* presents 4-6% above-ambient RH in its headspace, which decays to ambient levels within 30 mins after anthesis ^32^. Floral manipulations show that nectar evaporation accounts for half of floral humidity in *O. cespitosa*, while the other half results from petal transpiration. More recently, Harrap et al. ^33^ surveyed floral humidity from 42 plant species in a common garden, reporting a range of 0.05-3.7% above-ambient RH (henceforth ΔRH) in floral headspace. In lab assay, the hawkmoth *Hyles lineata*, a common pollinator of *O. cespitosa*, prefers probing unrewarded artificial flowers with above-ambient RH over flowers with ambient RH ^32^. Generalist pollinators like bumblebees (*Bombus terrestris*) can associate both dry and humid flowers with nectar presence under varied levels of background humidity, thus highlighting their flexible valence to floral humidity ^34^. These results indicate that floral humidity is informative to pollinators, influencing their foraging decisions while suggesting that our view of floral RH as a biophysical consequence of nectar evaporation may be overly simplistic.

Harrap and Rands ^35^ followed up their garden survey with manipulative experiments on two of their most humid flowers: *Calystegia sylvatica* and *Escholtzia californica*, yielding two important insights. First, funnel-shaped morning glory flowers produced the highest ΔRH values (∼3.7%) indicating that floral shapes that enclose the headspace can retain higher humidity levels^31^. Second, poppy flowers that offer pollen, but no nectar still produce >3% ΔRH, suggesting cuticular transpiration and stomatal conductance through the petals. These findings challenge the notion that floral RH is a cue that indicates floral profitability to pollinators while serving no adaptive function for flowering plants. The question of whether floral humidity may also function as a signal (like floral color and scent) hinges upon whether RH enhances floral fitness ^36,37^. In turn, this ultimate question, translated through pollinator service, depends upon several proximate issues, including physiological sources (and potential costs) of floral RH, the mechanisms of pollinator perceptual and behavioral responses to realistic RH gradients in space and time, and the efficacy of floral RH in the face of environmental noise.

We address these outstanding questions using the *Datura wrightii-Manduca sexta* plant-pollinator mutualism as a model system ^38,39^. Datura flowers present unusually high floral humidity (>30% ΔRH), exceeding levels previously reported for angiosperm flowers by a factor of 10. We show that Datura floral humidity is not a passive consequence of nectar evaporation and therefore is not an ephemeral cue. Instead, floral humidity in Datura is an outcome of gas exchange through floral stomates, thus is under the active physiological control of the plant. Neurophysiological responses from Manduca antennal hygrosensing neurons confirm that they perceive minute differences in floral humidity and experimental occlusion of the hygrosensing sensillum abolishes their innate behavioral preference for humid flowers. Furthermore, their attraction towards humid flowers cannot be abolished by experimentally rewarding only the ambient flowers, indicating a receiver-bias towards floral RH. In summary, using a neuroethological approach combining floral physiology, animal behavior, and sensory physiology, we provide compelling evidence that floral humidity in Datura flowers functions as a signal, rather than a cue, in this nocturnal pollination system.

## Results

### Vertical gradient of Datura floral humidity

Flowers of *Datura wrightii* exhibit an appreciable vertical humidity gradient (ΔRH) within the floral tube, with greatest ΔRH at the base of the flower tube (= 0 mm) (mean ± SEM, 31.01±1.2 %, n=38) across a broad range of background ambient RH (Fig. 1A). At the opening of the flower, ∼ 70mm above the corolla base (midpoint of the transect), the floral humidity was 4.08±0.50% ΔRH (Fig. 1A). Horizontal transects taken at the flower opening showed 3.89 ± 0.42 % ΔRH at the mid-point, consistent with the ΔRH recorded for the vertical transect at the same location (Fig. S1A). At 140mm above the flower tube, floral humidity was only marginally higher (0.46±0.16% ΔRH) than the background. Within the floral tube, the highest ΔRH (38.82±2.90%; n=4) was recorded when ambient RH was 10-20%, while the lowest ΔRH (24.16 ± 0.76%; n=14) was observed when ambient RH was 50-60%. We compared the floral humidity curves across background RH levels using two model parameters: decay rate (*α*) of the curve and the intercept *y0* (see Methods). Multiple comparisons suggested that the intercept (*y0*) differs across the range of background humidity but that the decay rate (*α*) does not (Fig. S2 & Tables S1, S2). Specifically, floral humidity measured when background RH was 50-60% was much lower than when background RH was 10-20% (*t*=3.92, *P*=0.001), 20-30% (*t*=3.50, *P*=0.004), and 30-40% (*t*=4.32, *P*=0.0001). No other differences were found for floral humidity at any other levels of background RH.

**Figure 1.**
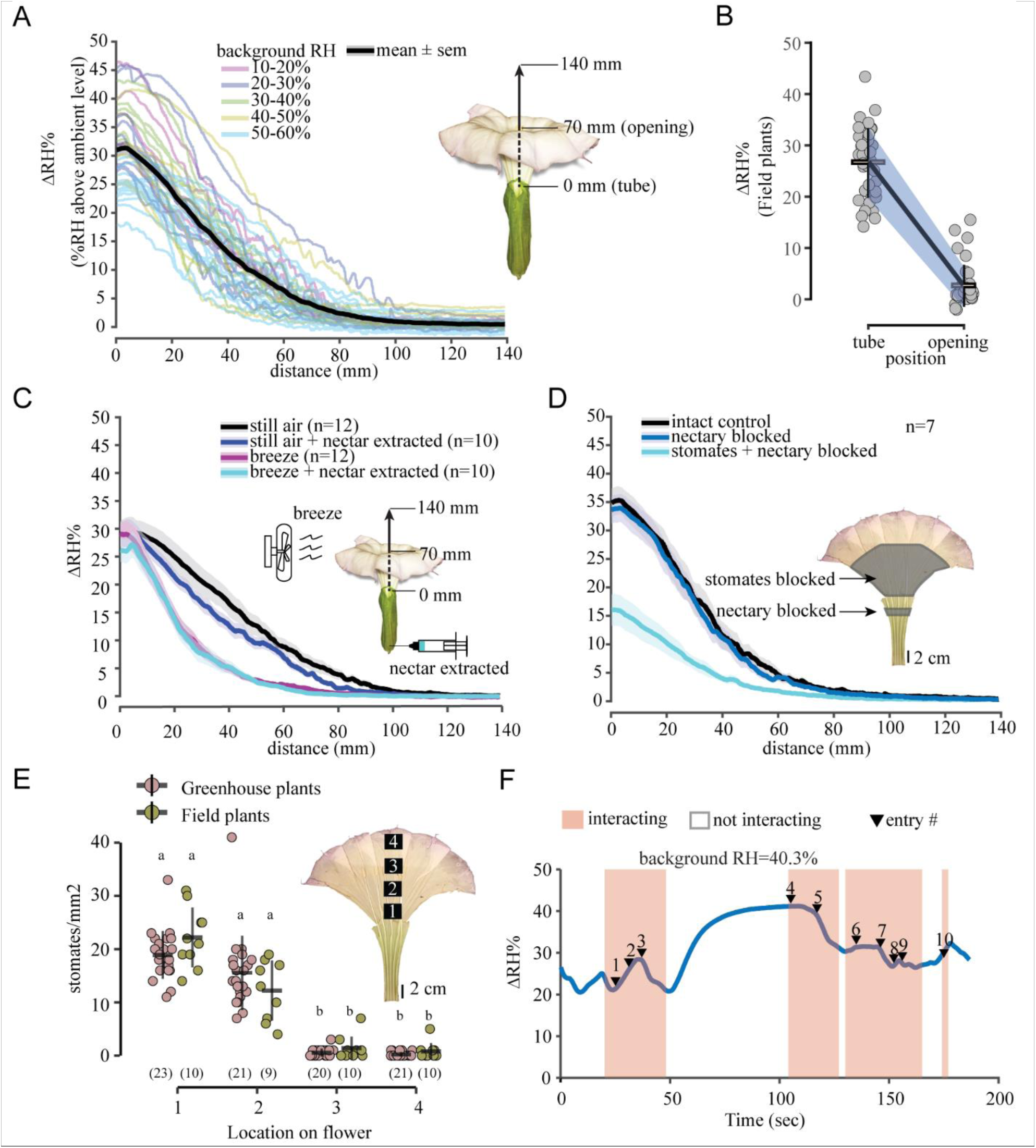
The efficacy of floral humidity as a cue for foraging pollinators. (A) A summary of the multiple transects of floral humidity measured from the base of the flower tube (0 mm) to outside the flower opening (140 mm) in a range of background humidity (10%-60% RH). See inset illustration of the vertical transect. Individual flower transects are color-coded by the background humidity they were measured at. Solid black line shows the mean, and the gray shading indicates ± SEM of *n=38* individual flowers. (Some transects are pooled from several experiments that are shown below). (B) Floral humidity of *n=37* naturally growing *Datura* flowers from Tucson, Arizona, USA, measured at two positions: flower opening (70 mm) and at the tube base (0 mm). Gray dots show individual flowers and line plots show the mean and standard deviation (blue shaded area). (C) Comparing the effect of breeze (∼ 0.4 m/s) and/or nectar extraction on the floral humidity. The inset figure illustrates the method for the different treatments. Treatments are color-coded showing mean (bold lines) and SEM (shaded area) with sample sizes in parentheses. (D) The effect of nectary blockage and stomatal blockage, as illustrated (inset), on the floral humidity curves. The mean (solid lines) ± SEM (shaded) are color-coded by treatment (*n=7* for each treatment). (E) Stomatal counts across 4 locations on the inner surface of the flower (see inset diagram) on greenhouse-grown plants (magenta) and field plants from Tucson, Arizona, USA (olive green). Dot plots show counts from individual flowers, line plots show the mean and standard deviation with sample sizes in parentheses. Letters show pairwise comparisons using the Wilcoxon test with Bonferroni correction. (F) Exemplary trace of floral humidity (solid blue line) measured continuously in the flower tube while moths interact with the flowers (orange shading) or enter inside the flower tube multiple times while probing (black triangles with the entry number). Floral humidity reconstitutes within seconds when the moth is not interacting with the flower (unshaded portions).

To confirm that the unusually high floral humidity in Datura is not an artifact of greenhouse conditions, we sampled Datura flowers in their natural habitat at multiple locations near Tucson, Pima Co., Arizona, USA (Aug. 2019; see Methods for site details). Even under field conditions and at backgrounds of 20-40% RH we recorded 26.58±6.71% (mean±SD) ΔRH in the flower tube and 2.64±4.02% ΔRH at the flower opening (Fig. 1B & Table S3).

### Efficacy of floral humidity

We performed a series of floral manipulations to test the efficacy of floral humidity under natural settings. We sampled floral humidity in still air as a control and subsequently added a gentle breeze to evaluate its effect on floral humidity (Fig 1C). Compared with flowers in still air, experimental breeze attenuated the vertical gradient of floral humidity (*α*:*t*= -6.6, *P*<0.0001; Table S4), but had no impact within the floral tube (*y0*: *t*=0.7, *P*=0.88; Table S5). We then extracted floral nectar to simulate moths probing and emptying the flowers. We found no evidence that nectar depletion influences floral humidity when comparing flowers sampled in still air with or without nectar (Fig. 1C). The decay rate (*α*) and the intercept (*y0*) of the floral humidity gradient were statistically indistinguishable between the still air and still air + nectar extracted treatments (*α*: *t*= -1.4, *P*=0.48; *y0*: *t*=0.5, *P*=0.9; Table S4, S5). Finally, we subjected flowers to both breeze and nectar depletion but saw no difference in the *α* or the *y0* in comparison to the flowers in the breeze with nectar present condition (*α*: *t*=0.08, *P*= 0.99; *y0*: *t*=0.5, *P*=0.9; Table S4&S5). Overall, these results indicate that the presence of floral nectar is not sufficient to generate the observed floral RH gradients.

Next, to test whether wing fanning and entrance of a hovering moth into a Datura flower could deplete floral humidity, we allowed moths to forage on newly opened Datura flowers while simultaneously recording the humidity in the floral tube (supp video 1). Remarkably, floral humidity never decayed to ambient levels even as moths hovered at the flower opening (interact) or entered flowers while probing (entry#) with floral humidity reconstituting within 30 sec of moths leaving the flowers (Fig. 1F).

### Source(s) of floral humidity

If not nectar, what is the primary source of humidity in Datura flowers? We conducted a separate experiment to evaluate the relative contributions of nectar evaporation and floral transpiration to floral humidity in Datura. We first blocked the nectary with petroleum jelly and compared the floral humidity transect with unmanipulated flowers. As expected, nectary blockage did not impact floral humidity curves (Fig. 1D). The decay rate and the intercept of the floral humidity were identical for the control and nectary blocked flowers, as noted from the fitted model predictions (*α*:*t*= -0.1, *P*=0.9; *y0*: *t*= 0.2, *P*=0.9; Fig. S4, Tables S6, S7). However, when both the nectary and the inner corolla surface were blocked with petroleum jelly, the magnitude of ΔRH was halved (Fig. 1D). The humidity curves of the flowers with their nectary and stomates blocked showed a significantly smaller *y0* than the other two treatments (*y0*: t= 5.6, *P*<0.0001; & *y0*: t= 5.3, *P*<0.0001; Table S7), but the *α* did not differ significantly (*α*: t= -0.5, *P*=0.8; & *α*: t= -0.4, *P*=0.9; Table S6). These results, combined with the nectar removal experiments above, demonstrate that transpiration is likely the major source of floral humidity in *Datura wrightii*.

We hypothesized that Datura flowers may contain stomates on the corolla to facilitate water vapor emission. Floral peels across four locations from the tube base to the inner (adaxial) corolla limb (Fig. 1E inset) indicated that stomates were found within the corolla (Fig. S5) at high density near the tube base but were scarce to absent towards the flower limb (Fig. 1E). This pattern was consistent between flowers of greenhouse-grown and wild plants. Stomates were observed across all four zones sampled on the outer (abaxial) corolla surface, and mean stomatal density was greater for wild plants than for greenhouse-grown plants (Fig. S1B). These data support the hypothesis that physiological gas exchange, rather than nectar evaporation, is responsible for the steep floral humidity gradients we have measured in the laboratory and the field.

### Hygrosensation in pollinators

#### Anatomy of the hygrosensing sensillum

Previous anatomical surveys identified at least 2 classes of aporous putative hygro-thermosensory sensilla on both male and female antennae of Manduca ^40,41^. The coeloconic type B is a small (2µm) peg-in-pit sensillum not easily visualized with microscopy. In contrast, the styliform complex is a large (30-40 µm), flexible-peg type sensillum on the leading edge of each antennal annulus and is easily distinguishable from other sensory pegs (Fig. 2A[I]). The styliform complex is the largest sensillum on female Manduca antennae, whereas it is surrounded by many large (pheromone-detecting) trichoid sensilla on male Manduca antennae (Fig.2A[II], also see ^42^). The tip of each sensillum houses 3-5 papillae (Fig. 2A[III]) of 2 µm diameter each. Individual papilla house 3 dendrites; 2 cylindrical and 1 lamellate type, as one unit enclosed within a dendritic sheath, typical of hygro-thermo sensory function (see ^41,43^), Thus, 9-15 dendrites innervate each putative humidity sensing organ, depending on the number of papillae at the tip of the organ, repeated over ∼80 annuli in both antennae (Fig. 2B).

**Figure 2.**
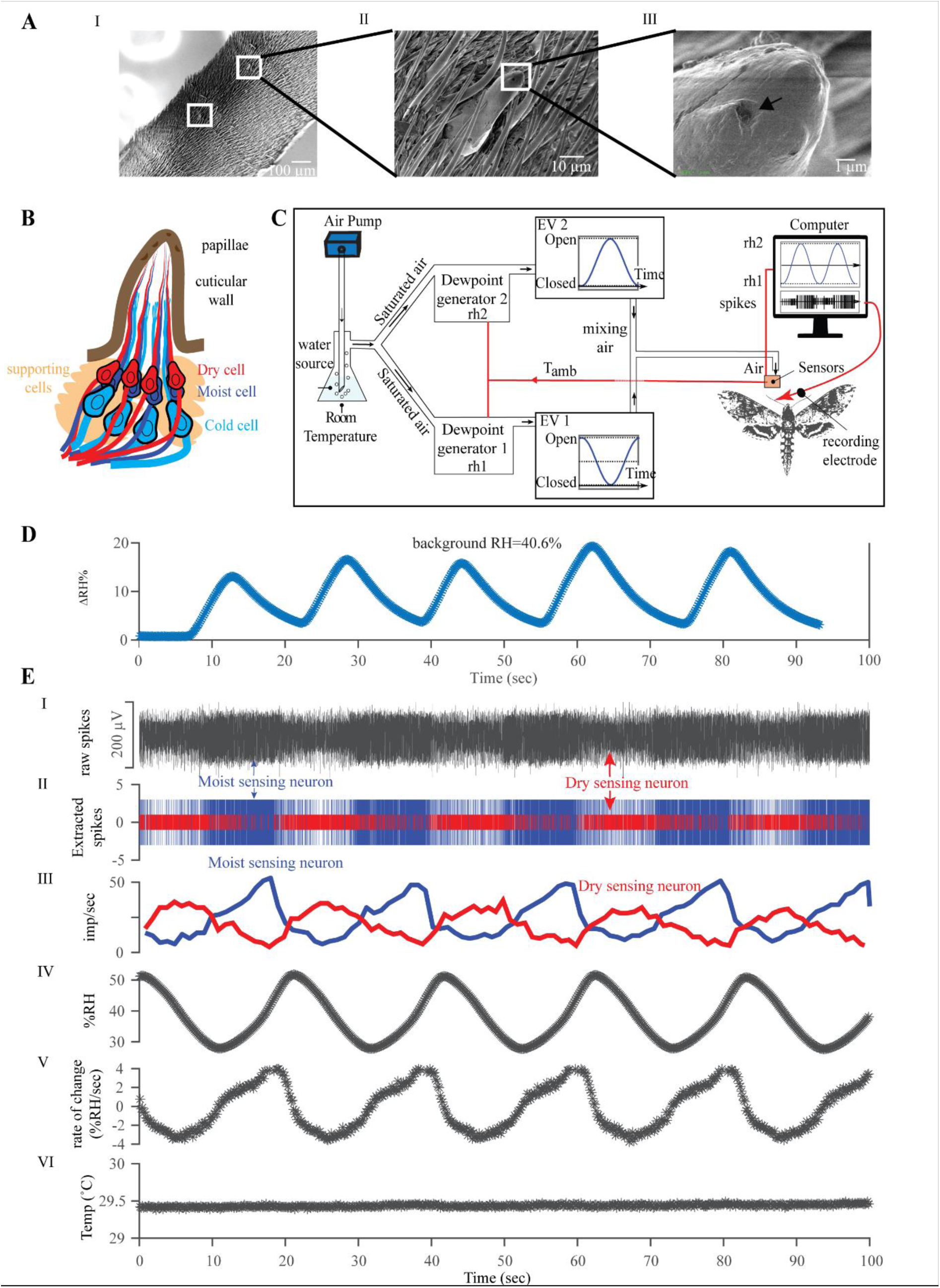
Moth antennal hygrosensory neurons respond robustly to the range of floral humidity presented by Datura flowers. (A) A zoomed sequence of SEM images of the styliform complex sensillum on *Manduca sexta* antenna. Scale bar is shown at the bottom right corner of each image. (I) Two segments of a female antenna with white squares showing the location of the styliform complex sensilla on the leading edge of the antenna. (II) A zoomed view of the entire styliform sensillum surrounded by trichoid sensilla. (III) Zoomed view of the tip of the styliform sensillum. Arrow indicates one of the papillae. (B) A representation of the longitudinal section of the styliform sensillum showing the underlying dendrites and cell bodies based on TEM ^40,41^ and cryosections of the organ. (C) Schematic of the stimulus delivery setup. Water vapor saturated air at room temperature is sent to two different dewpoint generators, outputting air with fixed relative humidity, *rh1* and *rh2*, corresponding to the ambient temperature, *T*_amb_ measured adjacent to the moth. Two electric valves (EV) operated by motors at the outlet of the dewpoint generators regulate the mass flow rate in an antiphase synchronized manner (as shown), which is sent to a thermostatic mixing valve to deliver air with a sinusoidally varying humidity airstream, like the fictive stimulus in (D). Temperature and RH were measured using sensors placed adjacent to the moth antennae. Tungsten electrode was poked at the sensillum base for electrophysiology. Electrical wirings are denoted in red, and black arrows denote the direction of airflow (see methods for details). (D) A fictive humidity stimulus generated by dipping the hygrosensing probe in a Datura flower to mimic the humidity experience of moths probing and entering Datura flowers. The blue line shows %ΔRH at background humidity of 40.6% (E) An exemplary single sensillum recording of the styliform sensillum (shown in A&B). (I) Simultaneously recorded extracellular activity of moist and dry neurons (arrows) within a single styliform sensillum. (II) An overlaid raster plot of the spikes sorted from the raw trace in (I) showing the activity of the moist sensing neuron (blue) and the dry sensing neuron (red). (III) A moving average of the impulse frequency of the moist (blue) and dry (red) neurons. (IV) A continuous sinusoidal stimulus of RH with amplitude ranging from 30% to 50% RH with a period of approximately 30sec. (V) Continuous rate of change in RH across the recording period. (VI) A constant temperature (ºC) is maintained across the recording period.

#### Fictive stimulus: floral humidity as experienced by flower-visiting moths

Simulating how moths probe Datura flower, the hygrosensing probe was dipped in and out of the flower to generate the range of humidity changes experienced by the hygrosensors on the moth antennae. Figure 2D shows the sinusoidal change in humidity as measured by the sensor entering and departing as a moth does while approaching and probing a Datura flower. At 40% background RH, the sensor measured a rapid increase of 15% ΔRH which dropped precipitously to ambient levels when removed from the flower.

#### Neurophysiology

We fashioned an experimental stimulus that temporally matches how moths might perceive RH as they enter and depart Datura flowers (Fig. 2D). The custom-built stimulus delivery system (Fig. 2C) generated a sinewave of RH whose amplitude and frequency could be altered to simulate the humidity change experienced by the hygrosensors on the moth antennae (Fig. 2E[IV]). Out of 39 electrophysiological recording events, 28 yielded responses to our stimulus from at least one type of sensory neuron, characterized as “moist”, “cold”, or “dry”. The underlying moist and dry sensing neurons (cold sensing neurons not evaluated here) responded robustly and predictably (Fig. 2E[I]) throughout our sinewave RH stimulus (Fig. 2E[IV]). This setup allowed us to maintain a stable temperature of the stimulus air while varying RH (Fig. 2E [VI]). The moist and dry neurons were distinguishable based on their amplitudes in most of the recordings (Fig. 2E[II]). The firing frequency of the moist neuron increased in proportion to the stimulus RH and remained well-correlated with the shape and phase of the rate of change of RH (Fig 2E[V]). The dry neuron increased firing frequency as the stimulus RH decreased and was ∼180 deg out of phase with the firing frequency of the moist neuron Fig. 2E[III]. Thus, the moist and dry sensing neurons showed the stereotypical antagonistic activity of the hygrosensory neurons previously demonstrated for other arthropods ^44-46^.

The few studied cases of insect hygrosensory neurons suggest that the response properties of the neurons are a function of both instantaneous humidity and the rate of change in humidity. Typically, the impulse frequencies of the moist sensing neuron increase in proportion to the instantaneous RH and rate of change in RH, whereas the dry sensing neurons respond antagonistically ^47,48^. To evaluate the response properties of Manduca hygrosensory neurons to these parameters, we fitted the data points with a polynomial linear regression of the form *F= a + b ΔRH/ΔT + c RH*, where *F* is the impulse frequency of the neuron, *a* is the height of the regression plane, *b* is the slope for the rate of change in RH, and *c* is the slope for the instantaneous RH. For both moist and dry sensing neurons, the slope for the rate of change in RH *b*, was larger than the slope for instantaneous RH, suggesting higher sensitivity to rate of change ±4% RH/sec, Fig. 2E[V] & Fig. 3A & B) compared to instantaneous RH (30-50%). For a rate of change of +1% RH/sec, this amounts to a decrease of -3.32 imp/sec for the dry sensing neuron, and correspondingly an increase of +4.24 imp/sec for the moist sensing neuron. For an increase in instantaneous RH by 1%, this amounts to a sensitivity of +0.28 imp/s for the dry sensing neuron, and -0.80 imp/sec for the moist sensing neuron. Calculations show that an increase of 1 imp/sec in the dry sensing neuron is elicited either by an increase of 3.56% instantaneous RH if the rate of change is constant, or by a rate of change of only -0.30%RH/sec. Similarly, for the moist sensing neuron, an increase of 1 imp/sec is reflected either by -1.24% instantaneous RH, if the rate of humidity change is constant, or by increasing the rate of change by only 0.23%RH/sec. Therefore, both hygrosensing neurons of Manduca are more influenced by the rate of humidity change of 1%RH/sec than by a 1% increase in instantaneous RH. This finding differs slightly from previous reports on other insects where the slope parameter *c* was found positive for the moist sensing neuron and negative for the dry sensing neuron ^47,49,50^. We conclude that Manduca’s styliform sensilla house hygrosensory neurons that are sensitive to minute fluctuations in humidity which moths likely experience as they hover at and enter floral headspace while probing.

**Figure 3.**
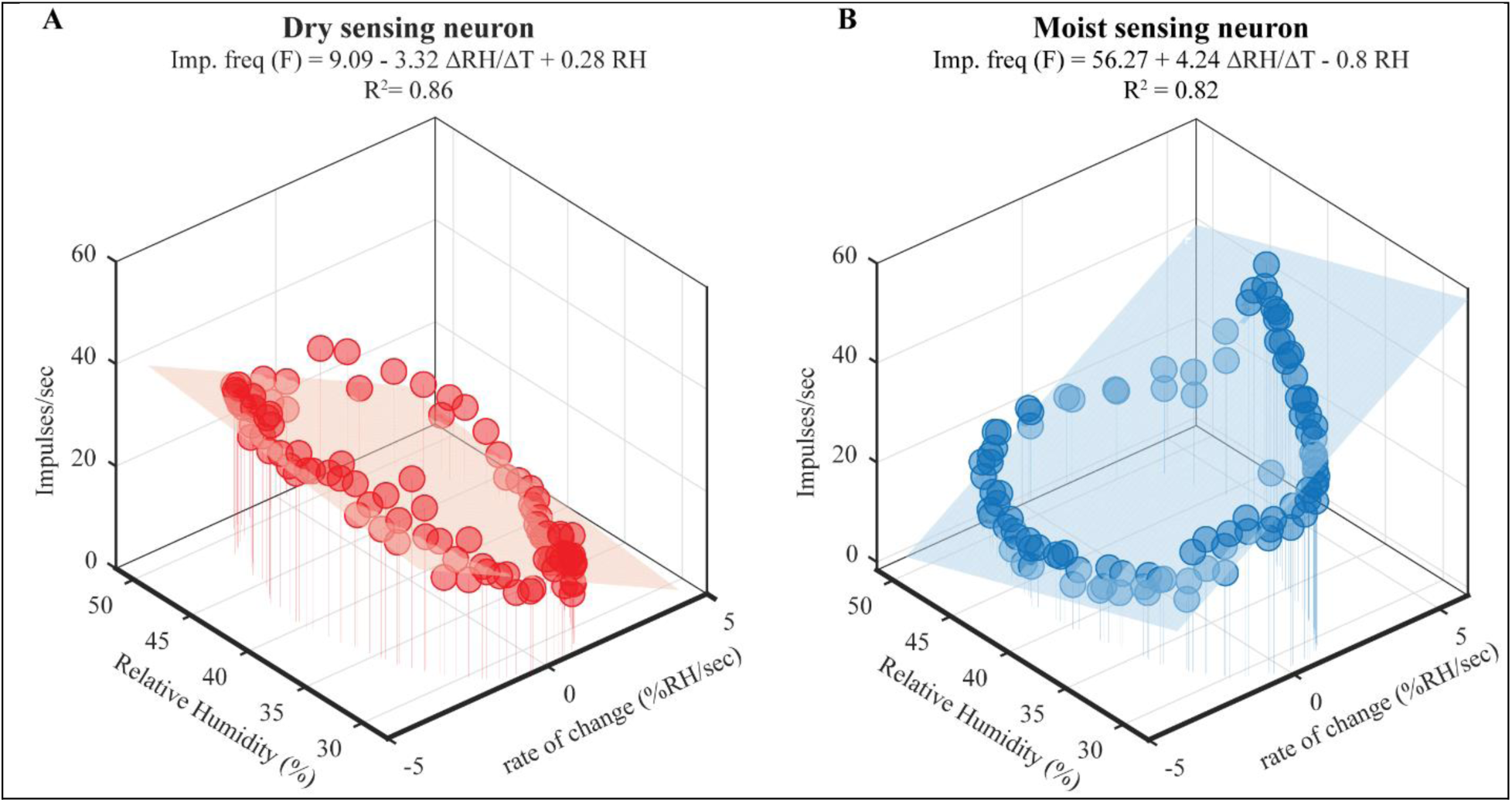
Response properties of the hygrosensory neurons are tuned to the rate of change in humidity. (A) & (B) 3D surface curve fitted scatterplots of the impulse frequency of the dry (A) and moist (B) neuron plotted against the rate of change in RH and the instantaneous RH. The fitted equation for the polynomial linear regression is shown at the top of each panel along with their goodness-of-fit measure (R^2^).

### Behavior

#### Flower-naïve hawkmoth preference for unrewarded ambient vs. humid artificial flowers

We presented flower-naïve adult Manduca with a choice between nectarless flowers with ambient humidity (henceforth, “ambient flowers”) vs. above-ambient humidity (henceforth, “humid flowers”) (Fig. 4A). The humid flowers presented a range of ΔRH matching the humidity measured from Datura flowers (Fig. 4B and S2). We used video tracking software to measure two response variables: probing duration and the number of floral entries made per visit. For the probing duration, we measured how long the proboscis tip (labeled) was within a circumscribed perimeter of the flower (region of interest shown in Fig. 3C[I]). Likewise, for the number of floral entries, we tracked how often the moth’s eye (labeled) crossed the flower rim (region of interest shown in Fig. 4D[I]). Neither males nor females showed side bias for probing duration (males: *Z*=0.29, *P*=0.76; females: *Z*=1.86, *P*=0.06; Fig. 4C[II]) or the number of entries (males: *Z*=0.39, *P*=0.68; females: *Z*=1.28, *P*=0.19; Fig. 4D[II]) when presented a choice between two identical ambient flowers. Irrespective of sex, moths probed longer on the humid flower than on the ambient flower (males: *Z*=3.85, *P*<0.001; females: *Z*=8.25, *P*<0.0001; Fig. 4C[III]) and entered humid flowers more frequently (males: *Z*=5.08, *P*<0.0001; females: *Z*=7.22, *P*<0.0001; Fig. 4D[III]). To assess if this strong preference for humid flowers is mediated through hygrosensation, we occluded a strip along the leading edge of the moth antennae with UV-hardened glue to block the styliform sensillum from contact with ambient air (Fig. S5B&D). Hygrosensor-blocked moths approached flowers less frequently as noted from the small number of visits shown in Fig. 4C&D[IV]. Nevertheless, the moths that did visit the flowers entered both ambient and humid flowers equally (males: *Z*=1.42, *P*=0.15; females: *Z*=0.003, *P*=0.99; Fig. 4D[IV]) and showed no strong preference for probing on either flower (males: *Z*=0.98, *P*=0.32; females: *Z*=0.62, *P*=0.53; Fig. 4C[IV]). Thus, moths with impaired antennal hygrosensation could not differentiate between humid and ambient flowers. To account for unintended effects of glue on moth antennae, we included a sham treatment where 5-10 annuli of the antennae were coated with the glue (Fig. S6B&D). Using racemic linalool as a common floral odorant, we evaluated the electroantennogram response of whole antennae of the hygrosensor blocked and sham control moths to test whether the antennae are still competent after the glue treatment. EAG responses showed that both treatments retain olfactory sensitivity to linalool (Fig. S6 A&C). Sham control moths retained a probing preference for the humid flowers (males: *Z*=4.04, *P*<0.0001; females: *Z*=3.43, *P*<0.001; Fig. 4C[V]) and entered humid flowers more frequently, just as unmanipulated moths do (males: *Z*=3.48, *P*<0.001; females: *Z*=3, *P*<0.01; Fig. 4D[V]). This confirmed that the glue occlusion was local and did not affect normal olfactory responses. In summary, both male and female flower-naïve Manduca show innate preferences for humid flowers. Their preference for empty humid flowers persists all night, despite the absence of nectar rewards.

**Figure 4.**
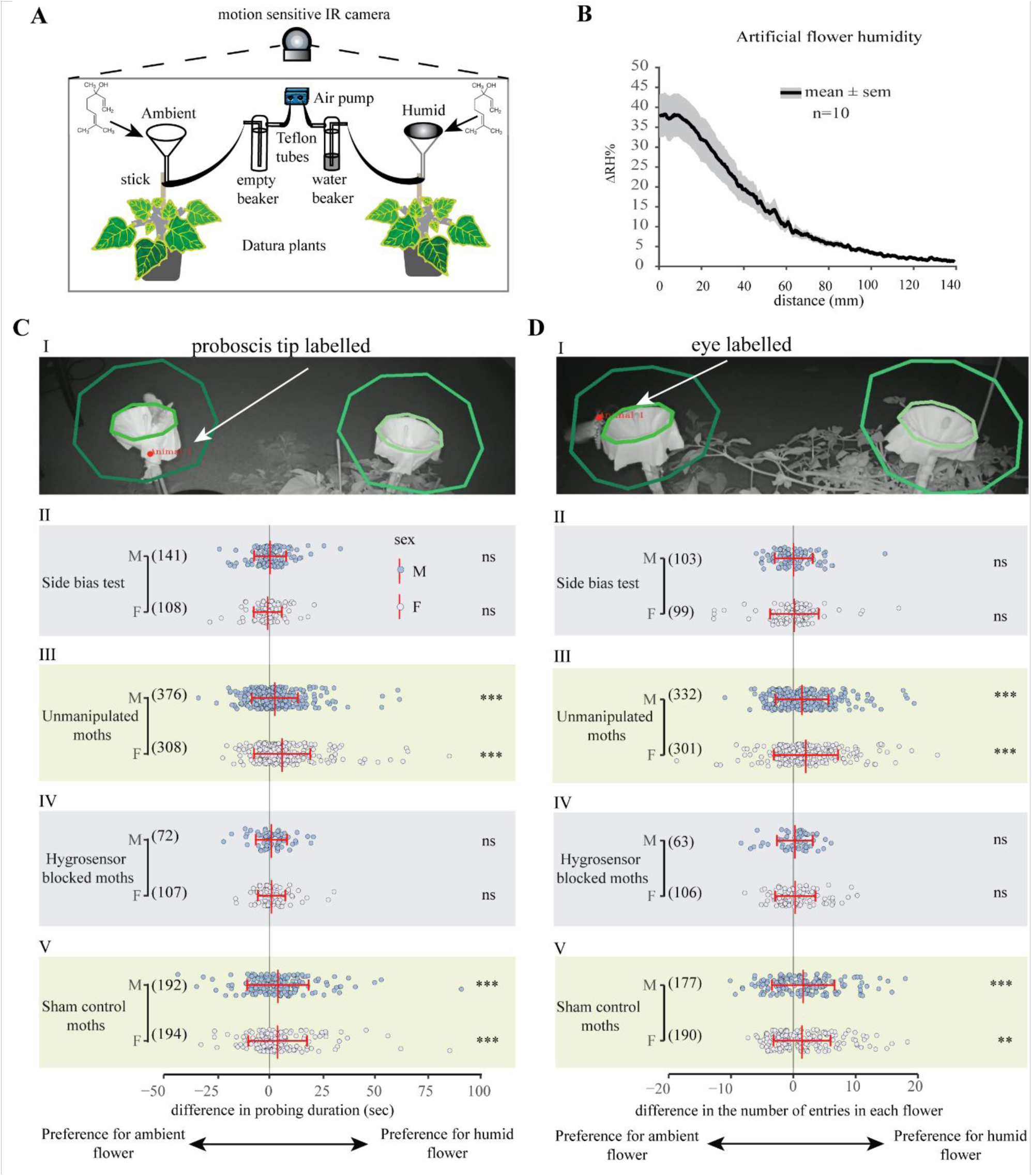
Moths are innately attracted to humid flowers in a binary-choice behavior assay. (A) Setup for the two-choice behavior assay using artificial flowers mounted on two non-flowering Datura plants. Chemical structures indicate the addition of scent (oil of bergamot) to both artificial flowers (funnels) at the start of the experiment. Air was pushed through the base of the funnels using an air pump (blue). For humid flowers, the air was pushed via Teflon tubes into a water beaker to generate saturated air, whereas, for ambient flowers, the air was pushed through an empty beaker. Overnight moth visits were video recorded using a motion-sensing IR camera (see methods for details). (B) Measurement of the vertical gradient of floral humidity of the artificial humid flower used in the behavioral experiment. Data are shown for n=10 transects as mean (solid line) ± SEM (gray shading). (C) & (D) [I-V] Behavioral responses of male (slate blue) and female (pale purple) naïve moths for the indicated treatments towards ambient and humid flowers, except for the side bias test in which both flowers presented ambient humidity. Differences in the duration of proboscis contact on each flower and the number of entries in each flower are the two measured response variables. Numbers in parentheses indicate the number of videos in which the labeled body part appeared in the region of interest for further analysis. The number of trials for each treatment was as follows: Side bias test: n= 12 nights for both sexes; Unmanipulated moths: n=11 nights for both sexes, Hygrosensor blocked moths: n=10 nights for both sexes; Sham control moths: n=12 nights for males, n=13 nights for females. For each night of the experiment, 2-4 naïve moths were released in the behavior room. Statistical differences are shown on the right side of each panel: ns, not significant, **p<0.01, ***p<0.001 (One-sample Wilcoxon signed-rank test against zero).

#### Moth preference for sugar-rewarded ambient and humid artificial flowers

To further understand the role of floral humidity as a signal or a cue for foraging hawkmoths, we modified the artificial flowers used in our binary choice assay to include sugar rewards (see Fig. S7B & C for flower design). We outlined three hypotheses decoupling the presence/absence of sugar rewards with floral humidity in accordance with key predictions related to floral profitability (Table 1).

**Table 1.** Hypothesis and predictions for experimental treatments decoupling floral humidity and sugar rewards. + indicates the addition of 25% sugar solution in the artificial flower, - indicates an empty flower.

| Flowers with or w/o rewards |  | Hypotheses |  | Predictions |
| --- | --- | --- | --- | --- |
| Ambient | Humid |  |  |  |
| + | + | Neutral cue | Because both humid and ambient flowers provide rewards, humidity is a neutral cue. | Weak but insignificant preference for the humid flower based on findings from the previous experiment (Fig. 5A[I]). |
| - | + | Index cue | Because only humid flowers provide rewards, humidity is an index cue for nectar presence. | Moths' innate preference for humid flowers is rewarded, resulting in a strong bias for the humid flower (Fig. 5A[II]). |
| + | - | Unreliable cue | Because humid flowers are empty and only ambient flowers provide rewards, floral humidity is an unreliable cue. | Moths learn to override their innate preference for the humid flower and strongly prefer visiting the rewarded ambient flower (Fig. 5A[III]). |

First, we tested the neutral cue hypothesis, where both ambient and humid flowers were provided with equal sugar rewards. We predicted that moths would show a weak, transient preference toward the humid flower over the ambient flower because floral humidity, in this case, does not provide additional information if both flowers are equally rewarding (Table. 1 and Fig. 5A[I]). Contrary to our prediction, moths showed a strong, persistent preference for the humid flower. Moths probed longer on the humid flower (*Z*=2.73, *P*<0.01; Fig. 5B[I]) and entered humid flowers more frequently (*t*=2.69, *P*<0.01; Fig. 5C[I]). To test the index cue hypothesis, we paired only the humid flower with the sugar reward but kept the ambient flower empty. We predicted that moths would show enhanced attraction to the humid flower (Table 1 and Fig. 5A[II]). The responses of moths in this experiment supported our hypothesis, suggesting that moths are quick to associate humidity with nectar presence. Moths probed and entered the humid flower with reward much more frequently than the empty ambient flower (probing duration: *t*=5.98, *P*<0.0001, Fig. 5B[II]; entries: *t*=4.6, *P*<0.0001, Fig. 5C[II]). Finally, we tested the unreliable cue hypothesis by adding a sugar reward to the ambient flower but keeping the humid flower empty, such that floral humidity is deemed unreliable for finding nectar. In this case, we predicted that the payoff disparity would lead moths to override their innate preference for the humid flower by visiting the rewarding ambient flower (Table 1 and Fig. 5A[III]). Contrary to our prediction, moths did not prefer the ambient flower with the sugar reward over the empty humid flower for either probing duration or the number of entries in the flower (probing duration: *Z*=0.75, *P*=0.45, Fig. 5B[III]; entries: *Z*=0.14, *P*=0.88, Fig. 5C[III]). Altogether, our results reject the hypotheses that floral humidity is a neutral cue or an unreliable cue. Our data are consistent with the predictions for floral humidity as an index cue, but this is insufficient because moths show perseverance to humid flowers even when reward is decoupled from floral humidity. Instead, moths show a strong ‘receiver-bias’ towards humidity, just as other insects demonstrate strong innate biases towards blue flowers, even when those of other colors are rewarding ^51,52^. This evidence, combined with our demonstration that nectar does not directly generate floral humidity gradients in Datura (Fig. 1), supports the role of floral humidity as a signal, not a cue, in this nocturnal plant-pollinator interaction.

**Figure 5.**
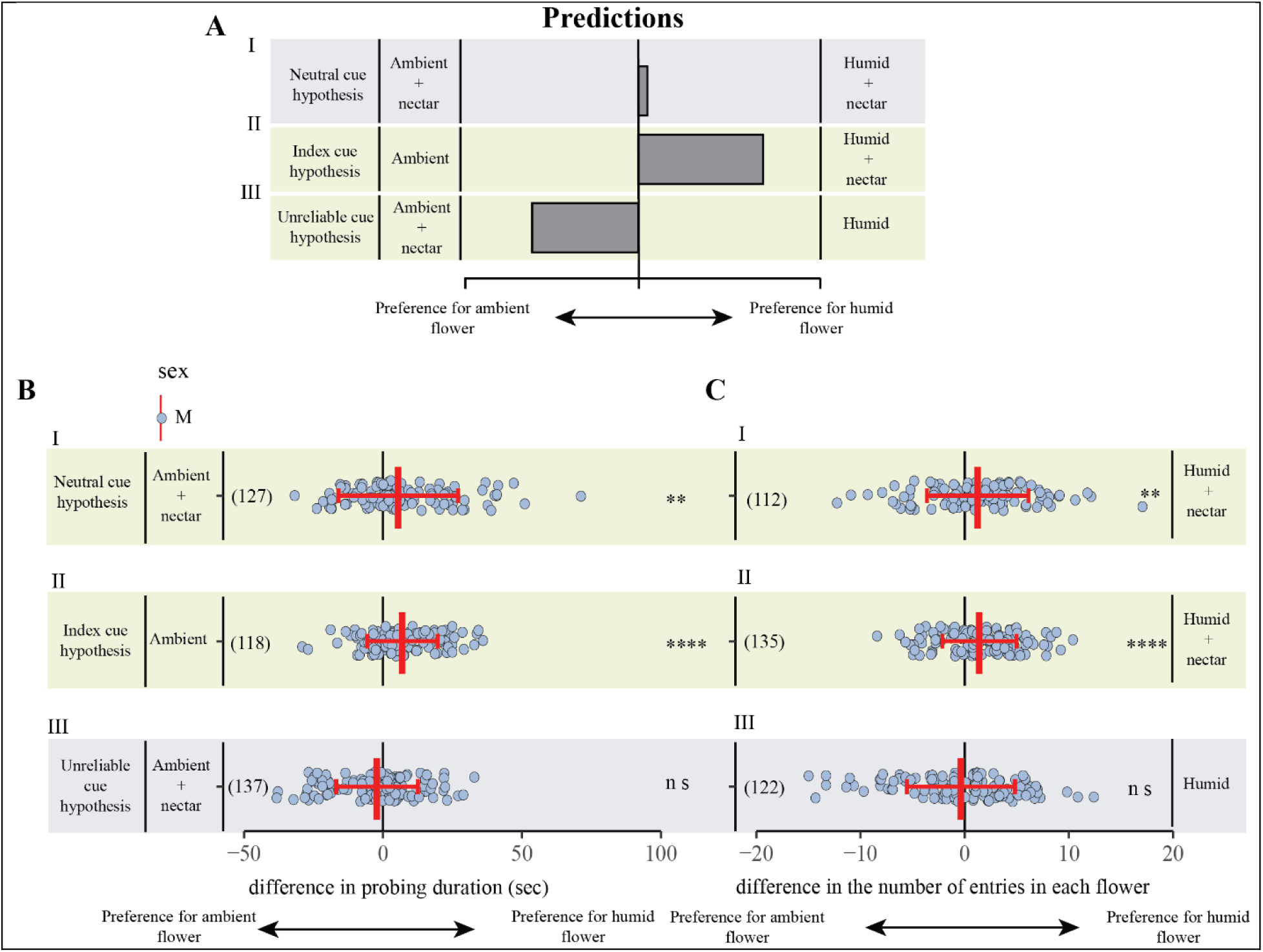
Moth responses to floral humidity decoupled with rewards reveal the signal-like role of floral humidity. (A) Hypotheses and qualitative predictions for moth responses towards humid and ambient flowers presented with sugar reward in either one or both flowers (see table. 1 for details). (I) Neutral cue hypothesis: both ambient and humid flowers provide nectar. (II) Index cue hypothesis: sugar reward in the humid flower, the ambient flower is empty. (III) Unreliable cue hypothesis: Ambient flower with a sugar reward, humid flower empty (B) & (C) [I-III] Experimental results of the three hypotheses outlined above. response variables are the same as in Fig. 3C & D. Data show individual visits (light blue dots) with mean ± SD (red lines) for each indicated hypothesis. Note that only males were used for this experiment. Numbers in parentheses indicate the number of videos in which the labeled body part appeared in the region of interest for further analysis. The number of trials for each treatment was as follows-neutral cue hypothesis: n=10 nights; index cue hypothesis: n=10 nights; unreliable cue hypothesis: n=9 nights. Statistical differences are shown on the right side of each panel: ns, not significant, **p<0.01, ****p<0.0001 (One-sample t-test or One-sample Wilcoxon signed-rank test against zero).

## Discussion

A key aspect of animal-plant communication is the distinction of traits into signals and cues. Cues could evolve into signals if they benefit both senders and receivers and incur a signaling cost ^25,26,53^. Senders must produce signals that are efficacious under most conditions and can be processed reliably by receivers ^26,54^. Current evidence for the role of floral humidity in pollination is scarce but suggests that it is a profitable cue for nectar presence ^20^. However, the evidence presented here reflects the role of floral humidity as a signal in a nocturnal pollination system. Through extensive floral manipulations we conclude that, unlike in *Oenothera cespitosa*, floral RH in Datura is not a consequence of nectar evaporation and therefore is not an ephemeral cue. We present evidence for the role of stomata in the production of floral humidity. Using an unbiased approach with automated tracking of moth body parts, we analyzed over 1200 videos of moths interacting with humid vs. ambient artificial flowers in overnight binary choice assays. We find that Manduca are innately attracted to humid flowers, a preference that is not limited to the first choice as reported for floral CO_2_ but persists for an entire evening of foraging. Using single sensillum electrophysiological recordings with a custom-built stimulus delivery setup, we show that Manduca antennal hygrosensing neurons are sensitive to the range of humidity offered by the flowers and can track temporal oscillations (e.g., resulting from breeze) in real-time. Furthermore, occluding the hygrosensing sensilla abolishes moths’ preference for humid flowers. The innate preference of moths towards humid flowers cannot be overridden when moths encounter flowers with ambient RH that are more rewarding, indicating a strong ‘receiver bias’ towards floral humidity. The additional probing that results from such bias would result in greater pollen export from Datura flowers^55^ potentially enhancing their male fitness through siring success and depositing more pollen on the stigma resulting in increased seed set ^38,56^. Below, we discuss the proximate and ultimate evidence supporting the interpretation that floral humidity functions as a signal in this pollination system.

### Proximate mechanisms

#### The efficacy of floral humidity as a stimulus

Despite a resurgence of interest in floral humidity ^32,33^, little is known about its efficacy to attract pollinators in the presence of background humidity, wind, and other physical factors. The humidity gradients (ΔRH) of Datura flowers greatly exceed background noise (plant or leaf headspace) or the calibration errors of most measuring probes (±2%RH) ^57^. Even during an exceptionally humid monsoon in Tucson, Arizona, USA (> 80% background RH in 2021), floral humidity remained 7-11% above ambient (Dahake personal observation). The maintenance of humidity gradients while moths visit Datura flowers underscores that floral humidity is a persistent physiological signature, rather than an ephemeral cue, in this system ^58^. In contrast, floral CO_2_ gradients, also likely established via corolla stomates, diminish after anthesis (with nectar secretion) in Datura flowers ^27^. In a different context, floral CO_2_ accompanies sustained respiratory activity in brood-site deceptive, thermogenic inflorescences of the dead horse arum (*Helicodiceros muscivorus*; ^59^) where it may fulfill the criteria of a signal if it contributes to the attraction of fly pollinators that utilize carrion as a larval host.

Unlike floral scent, humidity is likely a local stimulus for pollinators because it decays from the boundary layers of the floral tube to a background RH which can vary enormously (Fig.1A; also see ^32,33^). Such a local stimulus will be encountered by moths at a flower’s threshold while hovering. The influx of air generated by the vortices shed by the hovering wings of hawkmoths may drive headspace humidity out of the flower and over the moth antennae, as demonstrated for olfactory stimuli ^60,61^. Humidity is certainly experienced by moths upon entering flowers (Fig. 1F & Fig. 2D). The finding that humidity is not dissipated by wing fanning or air displacement suggests that it is produced continuously by floral tissues, much like scent. The humidity of Datura is also replenished rapidly after a pollinator’s visit (within 30 sec, Fig. 1F), ensuring its presence for subsequent visitors.

#### Pollinator perception of floral humidity through hygrosensation

Insect hygrosensing structures are typically located on the antennae ^62^. *Manduca sexta* antennae include at least two putative hygrosensing sensilla: the coeloconic type B (peg-in-pit) and the styliform complex sensillum (large peg, Fig. 2A)^40,41^. Electrophysiological recordings showed antagonistic activity of the dry and moist sensing neurons (Fig. 2E), a stereotypical feature of hygro-thermo sensing sensilla in other insects ^44-46,63,64^. Both moist and dry sensing neurons were more sensitive to the rate of change in RH than to instantaneous RH (Fig. 3A & B). Rates of change as low as -0.30% RH/sec and +0.23% RH/sec were sufficient to trigger spike frequency changes of 1 imp/sec in dry and moist sensing neurons, respectively, whereas, for instantaneous RH, the dry sensing neuron required -3.56% ΔRH and the moist sensing neuron required +1.24% ΔRH for a spike frequency change of 1 imp/sec in Manduca moths. This range of sensitivity values is comparable to those reported for cockroaches, stick insects ^49^, and honeybees ^50^ and supports the observed behavioral preferences of the smaller hawkmoth *Hyles lineata*, which could differentiate artificial flowers presenting 4-8% ΔRH ^32^, and the evidence that bumblebees (*Bombus terristris*) can identify humid flowers only 2% above ambient RH^34^. These findings reveal that Manduca hygrosensing neurons could respond to floral humidity as the moths enter floral headspace, even if flowers show a marginal ΔRH.

### Ultimate mechanisms

#### Sources of floral humidity and the potential costs of signaling

In a challenging environment, the mechanisms of floral humidity production imply physiological costs with potential fitness consequences to the plant. The presence of stomates on the corolla suggests that Datura flowers demand a constant supply of water to maintain turgidity in their desert-grassland habitat ^58,65^. Our knowledge of floral water budget and their attendant costs emerges from independent studies of very different kinds of flowers. For example, the bee-pollinated flowers of the alpine skypilot *Polemonium viscosum* lack stomata or trichomes yet lose enough water to incur a photosynthetic cost to the plant. To offset the cost, plants regulate flower size, producing smaller flowers during dry periods and larger flowers during humid periods ^65^. A similar trend is reported for self-pollinating flowers of *Leptosiphon bicolor*, where flower size correlates positively with moisture levels ^66^. Under dry conditions, the leaves of *L. bicolor* close their stomates to compensate for the water loss from flowers. In the giant semelparous succulent, *Agave desertii*, the entire plant accrues substantial costs during the climactic flowering event, when a majority of its water budget is allocated to the growing inflorescence ^67^. Although we have not calculated the flowering cost for *Datura wrightii*, their extraordinary flower size and blooming phenology suggest a significant cost to the plant. They grow along roadsides or desert washes where water flow during the monsoon rain season is abundant ^38^. Mass flowering only occurs during the monsoon rain season (July-September) in the Sonoran Desert ^68,69^. Flowers open only after sunset thus escaping the high desert heat during the day and minimizing evaporative loss. And lastly, plants have tuberous roots that store water and nutrients supplied when water is scant ^70^. Like *Leptosiphon, Datura wrightii* can self-pollinate, however, plants double their fruit and seed set when pollinated by *Manduca sexta* ^38^. Floral peels taken at multiple points during the night, or the following morning revealed that stomates are open throughout the night but close partially in the morning. These observations suggest that humid flowers likely incur costs to the plant, both in maintaining humidity gradients and floral turgidity through the night, in the xeric environment where Datura grows naturally. In the case of *Datura wrightii*, these costs may be especially high owing to the large surface area and volume (83.9 ±14.4cm^3^, n=10) of the flower and because a single plant can produce dozens of flowers in a given evening ^38^.

#### Significance of floral humidity for pollinators and sender-receiver conflict

The probing behavior of moths signifies an appetitive context. Even in the absence of a reward, moths showed perseverance by probing longer on the humid flower than on the ambient flower. Moths also entered the humid flower more frequently while probing. Flower entry shows commitment to nectar-feeding, increasing the chances of pollen deposition and export, by facilitating pollen loading on moth body parts ^55^. In the no-reward assays, moth responses clearly showed that they are innately attracted to humid flowers (Fig.4C & D). Furthermore, occlusion of the hygrosensing sensilla abolished moth preference for humid flowers (Fig. 4C[IV] & D[IV]), whereas sham control moths maintain a preference for humid flowers (Fig. 4C[V] & D[V]), confirming that the preference is mediated through hygrosensation. In all experiments, moth preference for humid flowers persisted beyond the first choice. This finding is unlike their preference towards floral CO_2_ which diminished to chance level after the first choice, in the absence of differential reward ^30^. This variable preference of moths towards floral primary metabolites is consistent with the temporal dynamics of CO_2_ and humidity in *Datura* flowers. CO_2_ in Datura flowers dissipates to background levels within the first two hours after anthesis ^27^, however floral humidity is consistently high throughout the evening (personal observation). Such multimodal interplay between primary and secondary metabolites in shaping pollinator behavior has been addressed in the cycad cones of *Macrozamia lucida* with their thrips pollinators ^71^. Thrips were neutral to CO_2_ but were repelled strongly by high temperature, humidity, and plant volatile production. Such studies reveal the temporal dynamics of signals and cues in conjunction with pollinator behavior, advancing our knowledge of plant-pollinator communication beyond floral advertisement.

A longstanding question within ecological communication is whether signals should always benefit both parties or is there room for sensory exploitation? ^72^. It is in the plant’s interest to maximize pollination services for minimum cost, whereas pollinators aim to maximize fitness (energy gain) per flower visit. To understand the conflict in Datura flowers between floral humidity and nectar, and its impact on moth foraging behavior, we addressed 3 alternate hypotheses by decoupling the presence of nectar with RH gradients (Fig. 5C&D). The response of moths towards the “unreliable cue” manipulations was most interesting. Here, the ambient flowers were rewarding, but the humid flower was empty, presenting a conflict between the moth’s innate bias and its expectation of a reward. Although we predicted that moths would learn to ignore the empty humid flower, moths visited both flowers equally (Fig. 5C[III] & D[III]), seemingly unable to abolish their strong preference for floral humidity. This evidence presents a potential avenue for plants to exploit pollinator perceptual bias ^37,73^. However, in such cases, floral humidity is likely cheaper for plants to produce than nectar. Such examples of sender-receiver conflict arising from sensory exploitation of pollinators are not uncommon in plant-pollinator interactions ^22,73,74^.

Our findings suggest multiple explanations for the significance of floral humidity to foraging pollinators. One possibility is that floral humidity serves as an inviting stimulus prompting hovering moths to enter the flower. This “microhabitat” hypothesis may be most relevant for the many smaller arthropods that utilize Datura flowers for mating, breeding, and protection ^75^. Another possibility is that floral humidity indicates either the presence or the location of nectar, influencing the rate of discovery. A similar role has been implied for visual nectar guides in bumblebee-pollinated flowers ^76^. Additionally, floral humidity may indicate larger nectar volumes and nectar quality ensuring that on average moths benefit from attending to this signal. Nevertheless, from the plant perspective, multiple entries of moths into Datura flowers increase pollen loading, promote siring success and cross-pollination ^55^. Indeed, reduced probing by *Manduca sexta* due to low nectar presence (and likely lower humidity) has been shown to lower seed set and impact plant fitness ^56^.

Finally, humidity may have evolved as a communication channel between plants and pollinators in the early stages of plant-pollinator diffuse coevolution. Hygroreception is mediated by Ionotropic receptor genes (IR) ^77-80^ which are conserved across arthropods and have a more basal origin compared to olfactory receptors ^81^. The fact that cycad cone humidity is a repellent signal for thrips pollinators in a basal gymnosperm lends credence to the notion that floral RH is a plesiomorphic component of plant-insect communication ^71^. Collectively, the high relative humidity of Datura flowers, its efficacy in the face of varying ambient conditions, and the salient behavioral and physiological responses that it evokes from moth pollinators reveal an unexpected signaling channel in plant-pollinator interactions, the benefits of which are tilted towards plant fitness.

## Methods

### *Manduca sexta* colony

We raised *Manduca sexta* from egg to adult in a laboratory walk-in growth chamber maintained at 24ºC and 50-60% RH with a 16:8 light: dark cycle. Caterpillars were fed a cornmeal-based artificial diet prepared in the lab. Late-stage caterpillars were transferred to individual cavities in wooden pupation blocks. After 7-10 days of pupation in the wooden blocks, pupae were transferred to the greenhouse to a moth breeding cage and left with a tomato plant for egg collection. Pupae used for the experiments were isolated from the lab breeding colony, separated by sex, and placed in 35 × 35 × 60cm (BioQuip) cages until ready for experiments.

### *Datura wrightii* plants

*Datura wrightii* seeds from Tucson, Arizona, USA were requested from the seed bank at Radboud University, Nijmegen, Netherlands (Accession number: 944750169). Seeds were soaked in water for 24 hours, followed by a rinse in 50/50 bleach water, and further soaked for 2 days in 0.1% Gibberellic acid. After soaking, seeds were nicked and placed on a wet filter paper in a petri dish until they germinated. Germinated seeds were sowed in 1-gallon plastic pots and placed in the greenhouse facility at Mudd Hall, Cornell University, under a 16:8 light: dark cycle. As necessary, plants were re-potted in a 3-gallon plastic pot, trimmed as needed, and regularly watered with 21-5-20 fertilizer (nitrogen-phosphate-potash). Plants continued flowering throughout the year.

### Vertical and horizontal gradients of floral humidity

For floral humidity measurements, flowering Datura plants were brought to a laboratory room from the greenhouse after anthesis. Floral humidity transects were carried out during the first 2 hours after anthesis at varying levels of background humidity throughout the year. The room temperature was 23±2ºC and background RH varied from 12% to 68%. Typically, the background humidity was high in summer months (40-60%RH) and low in winter months (10-30% RH). In preparation for measuring vertical or horizontal humidity gradients, flowers were held straight using bamboo sticks and metal wires. The Omega, Inc. hygrosensor probe (model 314A) was screw-fixed to a syringe pump (kdScientific model 100) to move the sensor gradually but continuously in either vertical or horizontal transects (see ^32^).The starting point for the vertical transects was the base of the flower tube, with the endpoint a few centimeters above the flower opening. At the start of the transect, the probe was lowered to the base of the corolla tube near the opening of the nectaries, ensuring that the probe head did not damage the anther filaments and the style. A vertical transect of 140 mm was carried out once for individual flowers with the probe moving at 0.21 mm/s. The slow speed ensured minimal mixing of air and allowed the hygrosensor to equilibrate. Horizontal transects were carried out 0.5cm above the surface of the corolla limb. As with the vertical transects, 14cm horizontal transects were taken at a fixed height over the open flower across its diameter. The reference probe was placed 10-20cm away from the flower at the same height as the flower opening. Output from the hygrosensors was compiled in the software provided by Omega. The humidity and temperature data were stored for every second of the transect, amounting to 650 points per transect. Delta RH was calculated by subtracting the ambient RH from floral RH, and the data were visualized in MATLAB 2019. Floral humidity data were collected as described here for all floral manipulations and artificial control flowers. We sampled floral headspace humidity in the tube and at the opening of Datura flowers growing in natural settings near Tucson, Pima Co, Arizona, USA. These settings included an experimental plot at Roger Road in urban Tucson, belonging to the Univ. of Arizona (32°16’41.3”N 110°56’18.5”W, 715m), a piñon-juniper-boulder habitat at the upper elevational limits of the plant’s distribution at Windy Point (32°22’07.0”N 110°43’00.8”W, 2013m) in the Santa Catalina Mountains, and in natural grassland habitat in the Santa Rita Experimental Range (31°47’01.5”N 110°49’32.3”W, 1322m). We measured the ambient humidity and temperature adjacent to the flower and noted the weather conditions at each location (Table S3).

### Floral manipulation experiments

#### Effect of breeze and nectar extraction on floral humidity

Conditions in nature are unequivocally more dynamic than in the laboratory. We expected wind to reduce the boundary layers of the flower surface. To test that, we generated an artificial breeze of approximately 0.4 m/s over the flower using a clip-on fan (15 cm diameter) in a laboratory setting. A black air filter pad 50 × 100 × 0.5 cm was placed between the fan and the flower to reduce airspeed and create a laminar flow. The distance between the fan and the flower was roughly 15-20 cm.

If the floral RH gradient is an outcome of passive nectar evaporation, we would expect nectar removal during hawkmoth visits to reduce floral RH ^32^. We extracted floral nectar by using a 1 ml disposable syringe to pierce through the base of the nectar tube and remove nectar from the 5 individual nectaries of each Datura flower before initiating the transect.

The experiments were carried out in a specific order. First, vertical transects were taken from control (unmanipulated) flowers in still air. Subsequently, a gentle breeze was applied to the flowers, and another transect was taken for the “breeze” treatment. Next, the nectar was extracted from the flowers, and the fan was turned off to measure the humidity of the nectar extracted flowers in still air. Lastly, the fan was turned on and another transect was taken for the “breeze+nectar extracted” treatment.

#### Floral nectary and stomate blockage experiments

Another way to test the contribution of nectar diffusion to floral RH is to occlude the floral nectar tubes (see ^32^). Accordingly, each of the five individual nectaries of the Datura flower was blocked with petroleum jelly applied locally using a narrow tube connected to a syringe filled with the jelly. An alternative model to produce floral RH gradients is active gas exchange through floral stomata (a physiological mechanism), rather than (or complementary to) diffusion from nectar (a physical mechanism). To block the stomates of Datura flowers, petroleum jelly was smeared on the inner (adaxial) surface of the corolla as shown in fig1D (also see ^35^). These experiments were carried out in the following order. First, vertical gradients of humidity were measured for the control (unmanipulated) flower. Next, the nectary was blocked, and another transect was taken. Finally, the inner surface of the flower was coated with jelly and a vertical transect was measured. In a separate experiment, floral humidity of the control flower was followed by the humidity of a flower coated with the jelly on the outer (abaxial) surface of the corolla as a sham control for the use of petroleum jelly and its potential interaction with water vapor and flower health. There was no indication of flower damage from using the jelly. This was confirmed by leaving jelly-coated flowers on the plant overnight and visually comparing them with unmanipulated flowers the following morning.

### Stomatal counts

To account for the large surface area of the Datura flowers, its corolla was divided into four zones from the base of the flower tube (location 1) to the lower limb (location 4; see Fig. 1E). The corolla surface was peeled off by hand at these four locations to expose the thin epidermis on the inner surface. Epidermal peels were stained with dilute safranin for 15 sec, rinsed in water, and mounted on a slide with a coverslip to visualize them at 20x under a Nikon Eclipse 80i compound microscope. Digital photos were taken of the prepared slides for peels at each location (Fig. S5). Subsequently, to note the scale of the image, a picture was taken of a reference slide with a 1mm grid engraved. Stomata were counted manually within a 1mm^2^ area drawn on the images using image J.

### Simultaneous measurements of floral humidity and moth interactions with Datura flowers

Fully opened flowers were excised from the plant, immediately placed in a conical beaker filled with water, and placed in a nylon mesh insect cage (BioQuip, Inc.; 71 × 71 × 122cm) within a laboratory room. Only male moths were used in this experiment and were isolated from the lab colony on the day of eclosion. Moths were trained to visit and handle Datura flowers at least one night before the experiment was conducted. The SHT31-D hygrosensor (Adafruit) was used for this experiment. The hygrosensor was connected to an Arduino Uno that was connected to a computer and operated through a custom-written Matlab code. The sensor was programmed to collect 10 data points per sec (upper limit) and was left running to collect data for 3 minutes while a moth was introduced to the insect cage. The background RH was noted at the start of every trial. Light intensity in the room was 0.1lx, measured using a light meter (Reed LX-1102). Moth visits were filmed using a Canon DSLR camera (EOS Rebel T6i) at 60 frames/sec for the 3-minute duration the sensor recorded floral humidity for each trial. The video data were aligned with the floral humidity data using a custom Matlab script (see S. video 1).

To measure the floral humidity experienced by moths as they enter and exit flowers, the hygrosensor was inserted in the flowers for 5-6 sec and removed for approximately 10 sec to mimic the behavior of moths when introduced to a cage with a single Datura flower, based on actual visits of moths in our experiment (data not shown).

### Behavior

#### Two choice assays on naïve moths presented with empty flowers

2-4 pupae of both sexes were isolated from the lab colony and placed in separate nylon mesh insect cages 40 × 40 × 60 cm (BioQuip, Inc.) in the greenhouse. We used 4-5-day old, starved moths for the experiment. Moths were released in the experimental lab room 3 × 6 m (width x length) at dusk. Two potted non-flowering Datura plants were placed in the center of the room with two white funnels (Büchner funnels, 9cm diameter) covered with a white paper towel attached to the plants, to mimic Datura flowers. The spectral reflectance of the paper towel matches closely with the authentic Datura flowers (Fig. S7A). A small night light was plugged in the wall opposite the plants, and window blinds were closed, yielding light intensity less than 0.01lx. The artificial flowers were attached to a bamboo stick and inserted within the Datura pots approximately 50 cm above the ground and 50cm apart from each other. Either humid or ambient air was supplied to the base of the artificial flowers through Teflon tubes connected to an air pump with two outlets (Topfin AIR 4000). The ambient flower received air passed through an empty beaker, whereas the humid flower received air pushed through a water-filled beaker resulting in a 0.3-0.4 m/sec airflow at both flower openings. A 2ml syringe plunger was inserted inside the tube of the artificial flower to mimic the grooves of the authentic Datura flowers. A cotton-tipped swab was glued to the center of the plunger to occupy the space taken up by the stamens and style in an authentic flower (Fig. S7B). Each evening, 5μl bergamot oil was added to the cotton swabs of both artificial flowers to provide a standardized, surrogate floral scent. A motion-sensitive IR video camera (Amcrest IP3M-941B) was placed to film overnight moth visits to both flowers. Videos were downloaded the following morning from the micro-SD card and were saved on a hard drive under appropriate treatment folders. The Datura plants were replaced and cycled through the 10 plants that were available in the greenhouse. Care was taken that plants had no blooming flowers during the trials. The positions of the ambient and humid flowers were alternated every night of the experiment.

#### Two choice rewarded assays

For the rewarded assays, we used only flower-naïve male moths to exclude the oviposition context associated with female moths. The artificial flower was modified by attaching four pipette tips at the edges of the syringe plunger to create 4 nectary grooves in the artificial flower (Fig. S7C). Individual pipette tips were filled with ∼50µl of 25% sucrose solution, amounting to 200µl in each flower (upper limit of nectar offered by the *D. wrightii* flowers in greenhouse conditions). Depending on the hypothesis to be tested, either one or both flowers were provided with a 25% sucrose reward. The following morning, nectary tubes were checked for consumption of sugar rewards, washed, and dried before using them for another trial. For behavioral response analysis, we only selected the videos after the moths found the nectary and contacted the sugar solution. It was crucial to exclude the exploratory flights where naïve moths probed on the flower limb or the flower stem to find the nectary. Starved moths showed a stereotypical behavior after they contact the nectary sugar solution with their proboscis after landing on the artificial flowers: they cease fluttering their wings and remain on the flowers with their proboscis extended in the floral tube for a prolonged duration to consume the reward. The response variable ‘probing duration’ accounts for the time spent by moths consuming the available nectar.

#### Occlusion of the hygrosensing sensillum and sham control

3-day old moths were cold-anesthetized in a -20ºC freezer for 10 mins. Once anesthetized, moths were viewed under a dissection microscope ventral side up and dorsal side placed over a cold metal block. A UV light-activated glue (Riverruns) was used to occlude the hygrosensors on the moth antennae. The glue bottle opening was attached with a 20-200µl pipette tip for localized application of the glue. The glue was applied along the leading edge of the entire antenna, completely coating the styliform sensilla. The glue was hardened under a handheld UV flashlight for 1-2mins. For sham control, only 5-10 segments at the base of each antenna were coated with the glue, leaving the rest intact. After the occlusion, moths were returned to the greenhouse for 1-2 days to recover from the handling stress before using them for the behavior experiment. The morning after the trial, moths were inspected under the microscope to evaluate the coverage of the glue on the antennae. In some cases, moths were able to remove the glue presumably while cleaning their antennae. Trials performed with such moths were excluded from further analysis.

#### Behavior video tracking and analysis

We used an animal-pose tracking software SLEAP ^82,83^ to track pre-selected body parts on the moth to collect information on moth position and behavioral choice in the two-choice behavioral assays. We selected the eye, head, proboscis tip, thorax, wingtips, and the abdominal tip of the moth’s body for tracking-based analyses on videos obtained by the motion-sensing camera (Fig. S8A). We labeled 1669 frames across 137 videos representing different behavioral trial sessions. A neural network was trained using SLEAP v1.1.5 installed on a PC equipped with a Geforce RTX 2080 Ti graphics card. The network was trained for 83 epochs or until the network loss value plateaued. We qualitatively assessed the network prediction accuracy from tracked behavioral videos prior to applying the network to track all remaining behavioral videos.

To derive metrics of the moth’s body positions around the artificial flowers, we used SimBA ^84^ to analyze the positional output data from SLEAP for all tracked body points. Five regions of interest were drawn across each video to enclose the cup of each artificial flower, the space surrounding the flowers, and the entire frame (Fig. S8B). For the probing duration, the proboscis tip labels were used, whereas, for the number of entries made into each region of interest, the eye was used for its proximity to the moth’s antenna and thus the hygrosensing sensilla. The output files were subsequently analyzed in R (v.4.1.1). Videos with tracking anomalies (<5%) were hand-corrected and the data were entered manually.

### Electrophysiology and imaging

#### Humidity stimulus delivery setup

We used two air pumps (Uniclife UL25 Air Pump) to continuously push air at a flow rate of 2L/min. The air was bubbled through an air stone immersed in water at room temperature. The resulting air saturated with water vapor served as input to two dewpoint generators (DG-4 DewPoint Generator, Sable Systems International). The dewpoint generators were set to operate in relative humidity controller mode, so that they outputted air at a fixed relative humidity of *rh1* = 11% and *rh2* = 90% corresponding to the temperature of the area adjacent to the moth antennae, *T*amb which was measured using a thermistor probe connected to the dewpoint generator. The airstream outlet of the dewpoint generator was fitted with a needle valve (Stainless steel High flow metering valve, Swagelok Inc.) with its head attached to a stepper motor (28BYJ-48 ULN2003 5V Stepper Motor) controlled using Arduino Uno (Arduino Inc.) and a stepper motor driver board (5V Stepper Motor ULN2003 Driver Board). We used a program written in Arduino IDE (open source) for regulating the air flow rate by controlling the valve opening position, motor speed, and motor acceleration. The valves were regulated such that the air outlets from the two dewpoint generators were anti-phase as shown (see Fig. 2D). The air streams mixed and passed through a T-junction connector and delivered locally with an air speed of 0.5-1.3 m/sec at the recording site on the moth antenna. We placed temperature and humidity sensors (AdaFruit SHT31-D) near the antennae (within less than ∼2 cm) to simultaneously measure the temperature and humidity of the delivered airstream. This apparatus allowed us to control the rate of the sinusoidal humidity stimulus as well as offer stationery or step-like stimuli. We partially programmed the humidity stimulus through Matlab and viewed it as a MATLAB figure simultaneously with the instantaneous electrophysiology output.

#### Single sensillum recordings and spike sorting

We immobilized 2–3-day old moths in a 15ml falcon tube. The falcon tube base was cut off just enough for the moth’s head and antennae to protrude. The head of the moth was prevented from moving by fixing it to the tube base with a collar of dental wax. The moth’s proboscis was extended and fixed to the tube with more dental wax to prevent proboscis movement from interfering with the electrodes. Moths were placed ventral side up under the microscope on a 10_×_10_×_2cm (l _×_ w _×_ h) plexiglass block. The antenna was adhered to the plexiglass with a hand putty (Blu Tack). The styliform sensillum was viewed under a microscope (WILD M3C, Heerbrugg, Switzerland) at 40x zoom objective attached with 1x magnifying lens and 20x eyepiece. For single sensillum electrophysiology, a sharp 2.5cm wax-coated tungsten microelectrode (MicroProbes) with 2.5MΩ resistance and a similar reference electrode was attached to a headstage (A-M systems, model 1800) fixed on a micromanipulator (Narishige). All electrical components of the electrophysiology rig were grounded to a wire in the room away from the rig. The wires attached to the hygrosensor placed next to the moth antenna were wrapped in aluminum foil to reduce electrical noise. The recording electrode was connected to a two-channel high impedance amplifier (A-M systems model 1800), the signal was bandpass filtered for 300 to 1000Hz, a notch filter was turned on and the signal was transferred to a data acquisition device (National Instruments, Inc., model USB-6211). The data acquisition device was connected to a computer and the signal was visualized using the open-source software Spike Hound v1.2 ^85^. The sampling rate was set at 20,000Hz and the individual recording sessions 3-6 min each was saved on the computer until further analysis. Selected raw electrophysiology traces were spike-sorted using an open-source Waveclus 3.0 toolbox ^86^. The stimulus and the spikes were aligned, analyzed, and visualized in MATLAB (R2019a) using a custom script.

The micromanipulator was advanced to insert the recording electrode at the base of the styliform sensillum. The reference electrode, attached to an electrode holder and controlled by another micromanipulator, was inserted a few segments toward the proximal end of the same antenna as that of the recording electrode. Electrophysiology was performed on moths of both sexes, and a new moth was used for every new recording event. *M. sexta* antennae consist of one styliform sensillum on each segment of the antenna, thus, during one recording event, we attempted multiple sensilla on 5-10 segments of the middle portion of the moth antenna. The styliform sensillum is a complex of 3-5 individual sensing organs (papillae) located at the tip of the peg ^43^. Therefore, a high density of cell bodies is present beneath the sensillum (see Fig. 2B). Many recording events pick up more than one unit of one cell type. A moist neuron was identified if it responded with increasing firing frequency when the stimulus humidity increased, whereas a dry sensing neuron was identified if it showed increased impulse frequency when the stimulus humidity decreased ^47^. Therefore, the moist and dry sensing neurons are antagonistic to each other in their responses to changes in humidity. These two neurons are associated with a third neuron, the cold cell ^87^, that responds with high impulse frequency when the air temperature decreases but ceases firing when the air temperature increases (data not shown). For every new recording event, the amplitudes of the dry and moist sensing neurons vary depending on where the tip of the electrode is in relation to the cell bodies of the neurons. However, the ratio of the amplitude stayed constant throughout the length of the recording. We were able to record anywhere from a few minutes to a couple of hours from the cells within a sensillum. Males and females showed identical responses.

#### Scanning electron microscope images

SEM images were taken of air-dried antennal samples of both sexes under a Zeiss electron microscope. Samples were sputter-coated with gold for 30sec and imaged at EHT between 0.3 to 1 kV and WD between 2.4 to 7.7 mm.

#### Statistics

To evaluate statistical differences among floral humidity curves, we used R v.4.1.1 ‘nlme’ package to fit nonlinear models to the data^88^. We started with a simple non-linear mixed effect model with no effect of the different treatments and no random effect of the individual flowers on the model parameters. However, adding the effect of the different treatments in the fixed effects and the random effect of individual flowers, significantly improved the model and lowered the AIC value. Our final fitted nonlinear mixed effect model is as follows:

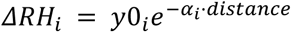

The best fitted model suggests that ΔRH (%RH above ambient) varies by treatment *i* and decays exponentially from the initial value *y*0 for that treatment, to the he final value at decay rate of *α* by distance. The model allows for separate intercepts and decay rate for each treatment *i* and includes random effects of individual flower transects on the intercept *y*0 and the decay rate *α*. Using package ‘emmeans’ we calculated the estimated marginal means (emm) and 95% confidence intervals for *y*0 and *α* for each treatment. We performed pairwise t-tests with post-hoc Tukey adjustments to the p values in comparing the *y*0 and *α* values between multiple treatments.

For the stomatal counts, we performed a Kruskal-Wallis test across the four locations at which we counted stomatal density. For all the behavior data, we performed either one-sample t-tests or Wilcoxon tests, depending on the distribution of the data (normal vs. not normal), with the null hypothesis being that the differences in probing duration and the number of entries between humid and ambient flowers are not different from zero. In other words, the null prediction is that moths cannot distinguish between humid and ambient flowers and visit both flowers equally.

We used MATLAB R2019b to generate the 3D scatterplots of cell impulse frequency (y-axis), plotted against and the instantaneous RH (x-axis), and rate of change of RH (z-axis). The MATLAB curve fitting app, cftool was used to fit three-dimensional polynomial linear regressions to the data of the form *F* = + *bΔRH/ΔT + cRH*, where *F* is the impulse frequency of the dry or the moist neuron, *a* is the height of the regression plane, *b* is the slope for the rate of change in RH, and c is the slope for instantaneous RH.

## Acknowledgements

We thank John Putnam for greenhouse care, Deidra Jacobsen for help germinating Datura seeds, and Cole Ortiz for help with stomatal counts. We are grateful to Ron Hoy and Gil Menda for providing the electrophysiology equipment. We thank Goggy Davidowitz for hosting A.D. and R.A.R in Tucson and for field assistance in measuring floral humidity. We are thankful to Gordon Smith and Shayla Salzman for their insightful comments on the earlier versions of the manuscript. Lynn Marie Johnson from the Cornell Statistical consulting unit provided excellent help on performing the non-linear models on floral humidity curves. This work made use of the Cornell Center for Materials Research Shared Facilities which are supported through the NSF MRSEC program (DMR-1719875). Parts of this research were funded through Cornell Sigma Xi, Cornell CALS alumni grant, Cornell neurobiology and behavior dept. grant awarded to A.D. The physiology data was part of the Cornell Neurotech Mong fellowship awarded to A.D. and P.J. in the labs of R.A.R. and A.S.

## Author contributions

A.D. and R.A.R. conceived the study, A.D. performed all experiments and analyzed data with help from P.J., C.V. and W.K., P.J. and A.S. designed the stimulus delivery setup. A.D. and R.A.R. wrote the paper; all authors edited the manuscript.

