## Supplementary information for "Floral humidity as a signal – not a cue – in a nocturnal pollination system"

**Supplementary material**

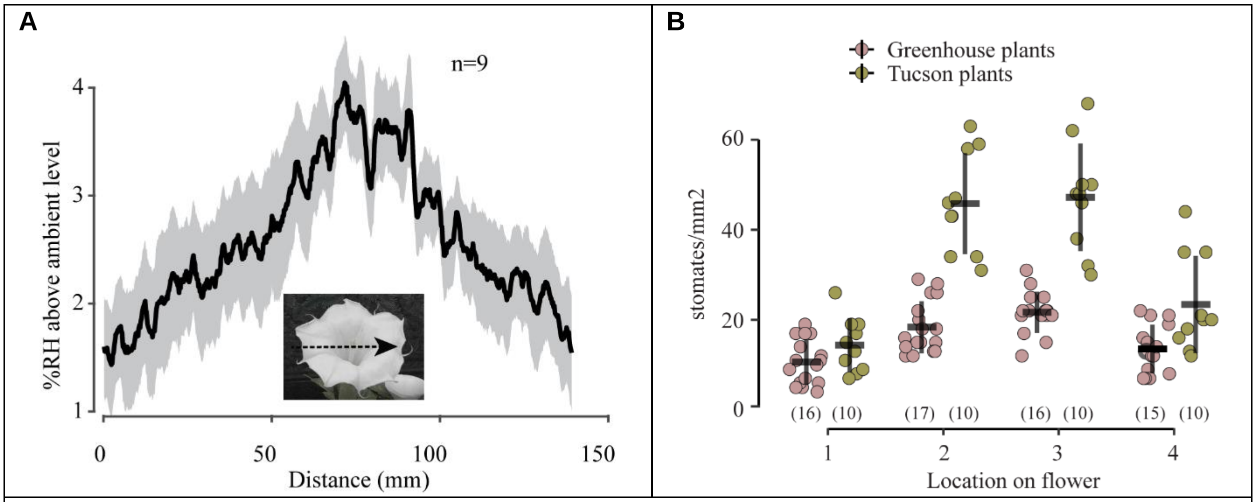


**Figure S1. (A)** Floral humidity measured along a horizontal transect 0.5cm above the flower opening. The inset image of a *Datura wrightii* flower with the arrow shows a typical transects trajectory. The sample size is indicated in the top-right corner.
**(B)** Stomatal counts across 4 locations, same as Fig. 1E, but on the outer surface of the flowers (abaxial) from greenhouse-grown plants (magenta) and flowers collected from Tucson, AZ, USA (olive green). Sample sizes are indicated in parentheses.

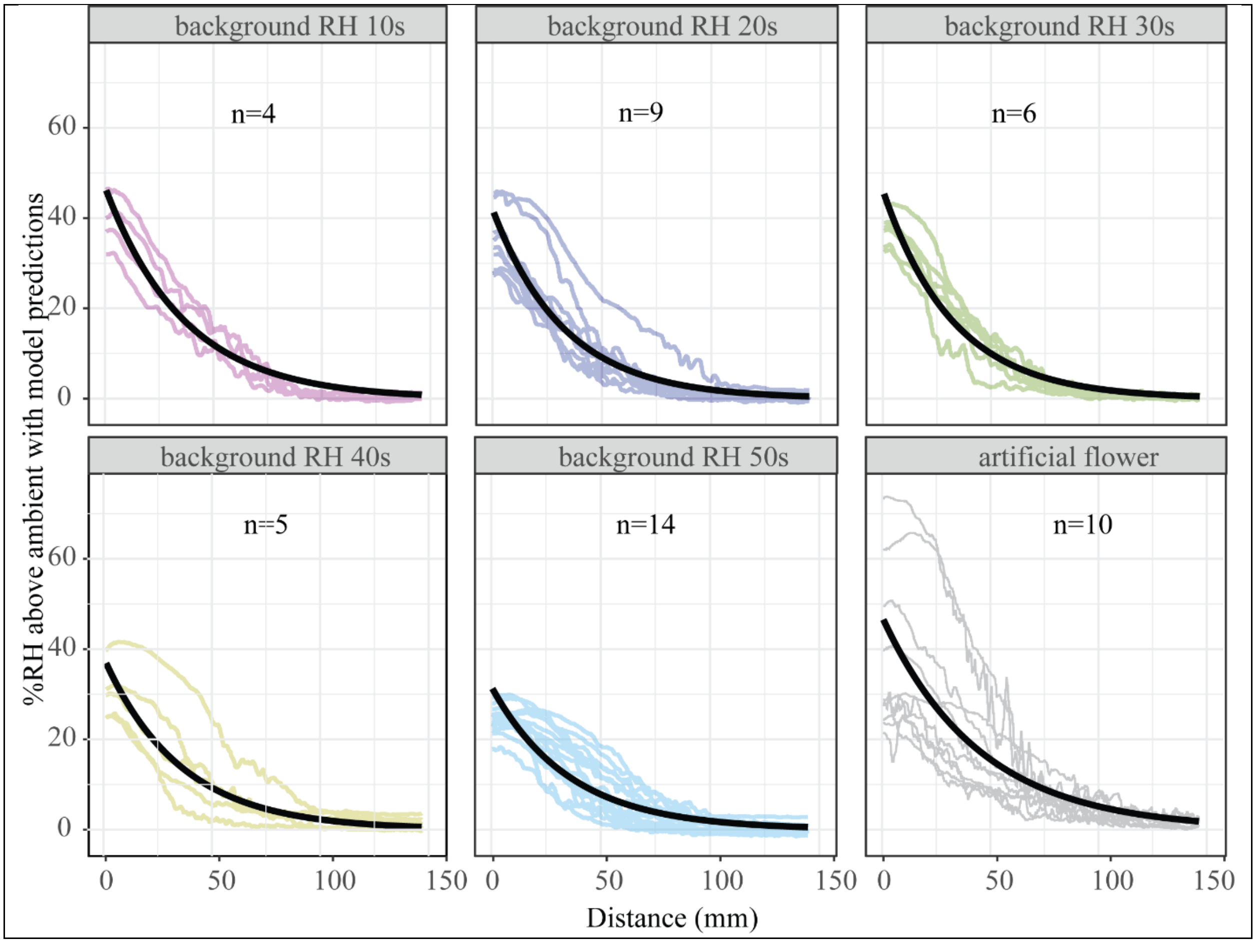

**Figure S2.** Predicted model fits (solid black line) for floral humidity at different levels of background humidity including the predicted fit for the humid artificial flower used in the behavioral experiments. Individual flower transects are shown for the indicated level of background humidity. Sample sizes are mentioned in each panel.

**Table S1.** Multiple comparisons of the model estimates for the **decay rate (α)** across the different levels of background humidity as shown in **figure S2** (post hoc Tukey HSD).

| comparisons | estimate | SE | df | T ratio | P-value |
|---|---|---|---|---|---|
| teens - twenties | -0.00303879 | 0.00419275 | 24653 | -0.72 | 0.95 |
| teens - thirties | -0.00353883 | 0.00450340 | 24653 | -0.79 | 0.93 |
| teens - forties | -0.00102431 | 0.00468254 | 24653 | -0.22 | 1.00 |
| teens - fifties | -0.00048512 | 0.00395634 | 24653 | -0.12 | 1.00 |
| twenties - thirties | -0.00050004 | 0.00367897 | 24653 | -0.14 | 1.00 |
| twenties - forties | 0.00201448 | 0.00389620 | 24653 | 0.52 | 0.99 |
| twenties - fifties | 0.00255367 | 0.00298443 | 24653 | 0.86 | 0.91 |
| thirties - forties | 0.00251451 | 0.00422870 | 24653 | 0.59 | 0.98 |
| thirties - fifties | 0.00305371 | 0.00340710 | 24653 | 0.90 | 0.90 |
| forties - fifties | 0.00053919 | 0.00364059 | 24653 | 0.15 | 1.00 |

**Table S2.** Multiple comparisons of the model estimates for the **intercept y0** across the different levels of background humidity as shown in **figure S2** (Post hoc Tukey HSD).

| comparisons | estimate | SE | df | T ratio | P-value |
|---|---|---|---|---|---|
| teens - twenties | 4.89352959 | 4.04871812 | 24653 | 1.209 | 0.746 |
| teens - thirties | 0.75496044 | 4.34911691 | 24653 | 0.174 | 1.000 |
| teens - forties | 9.30934183 | 4.51954395 | 24653 | 2.060 | 0.238 |
| teens - fifties | 14.9876402 | 3.81964569 | 24653 | 3.924 | 0.001 |
| twenties - thirties | -4.13856915 | 3.55136021 | 24653 | -1.165 | 0.771 |
| twenties - forties | 4.41581224 | 3.75814036 | 24653 | 1.175 | 0.766 |
| twenties - fifties | 10.0941106 | 2.878721 | 24653 | 3.506 | 0.004 |
| thirties - forties | 8.55438139 | 4.07999001 | 24653 | 2.097 | 0.221 |
| thirties - fifties | 14.2326798 | 3.287816 | 24653 | 4.329 | 0.0001 |
| forties - fifties | 5.6782984 | 3.5101558 | 24653 | 1.618 | 0.486 |

**Table S3.** Floral humidity of *Datura* flowers measured in their natural habitat during the week of Aug 14 to 19, 2019, at three locations in Pima county, Tucson, Arizona.

| location | Ambient conditions | | %ΔRH (mean ± SD) | | n | Weather conditions |
|---|---|---|---|---|---|---|
| | % RH | Temp ºC | tube | opening | | |
| University of AZ, experimental plot | 29 | 31.8 | 27.79±5.63 | 4.00±4.84 | 21 | breezy, moths probing flowers |
| Windy Point | 24 | 25.3 | 29.73±7.29 | 1.07±0.73 | 7 | windy, no moths |
| SRER, grassland | 33.9 | 28.7 | 21.33±5.62 | 0.69±1.13 | 9 | Overcast and strong gusts of wind, storm approaching. |

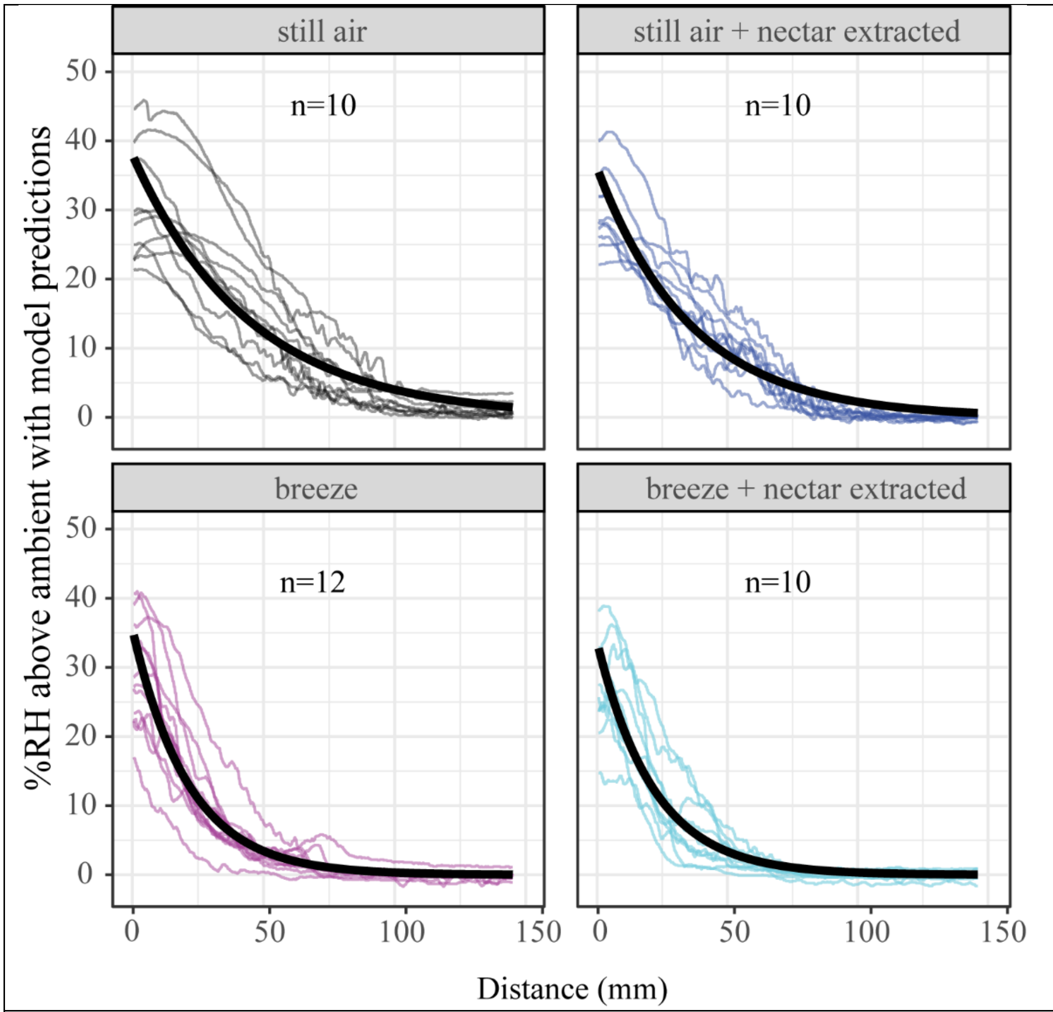


**Figure S3.** Model predictions (solid black line) for floral humidity were examined under different treatments. Individual flower transects (color-coded) are shown for the indicated treatment. Sample sizes are mentioned in each panel.

**Table S4.** Multiple comparisons of the model estimates for the **decay rate (α)** across the different floral manipulations shown in **figure S3** (Post hoc Tukey HSD).

| comparisons | estimate | SE | df | T ratio | P-value |
|---|---|---|---|---|---|
| control - nectar extracted | -0.0056754 | 0.00397353 | 27251 | -1.4283 | 0.4815598884 |
| control - breeze | -0.02521763 | 0.00381531 | 27251 | -6.6096 | 0.0000000002 |
| control - (breeze+nectar extracted) | -0.02489692 | 0.00398317 | 27251 | -6.2505 | 0.0000000025 |
| nectar extracted - breeze | -0.01954223 | 0.0038161 | 27251 | -5.1210 | 0.0000018152 |
| nectar extracted - (breeze+nectar extracted) | -0.01922152 | 0.00398393 | 27251 | -4.8248 | 0.0000083445 |
| breeze - (breeze+nectar extracted) | 0.00032071 | 0.00382614 | 27251 | 0.0838 | 0.9997890394 |

**Table S5**. Multiple comparisons of the model estimates for the **intercept y0** across the different floral manipulations shown in **figure S3** (Post hoc Tukey HSD).

| comparisons | estimate | SE | df | T ratio | P-value |
|---|---|---|---|---|---|
| control - nectar extracted | 2.04763868 | 3.853107 | 27251 | 0.531 | 0.951 |
| control - breeze | 2.67904836 | 3.68976469 | 27251 | 0.726 | 0.887 |
| control - (breeze+nectar extracted) | 4.63361166 | 3.85392027 | 27251 | 1.202 | 0.625 |
| nectar extracted - breeze | 0.63140968 | 3.69000085 | 27251 | 0.171 | 0.998 |
| nectar extracted - (breeze+nectar extracted) | 2.58597298 | 3.85414637 | 27251 | 0.671 | 0.908 |
| breeze - (breeze+nectar extracted) | 1.9545633 | 3.69085005 | 27251 | 0.530 | 0.952 |

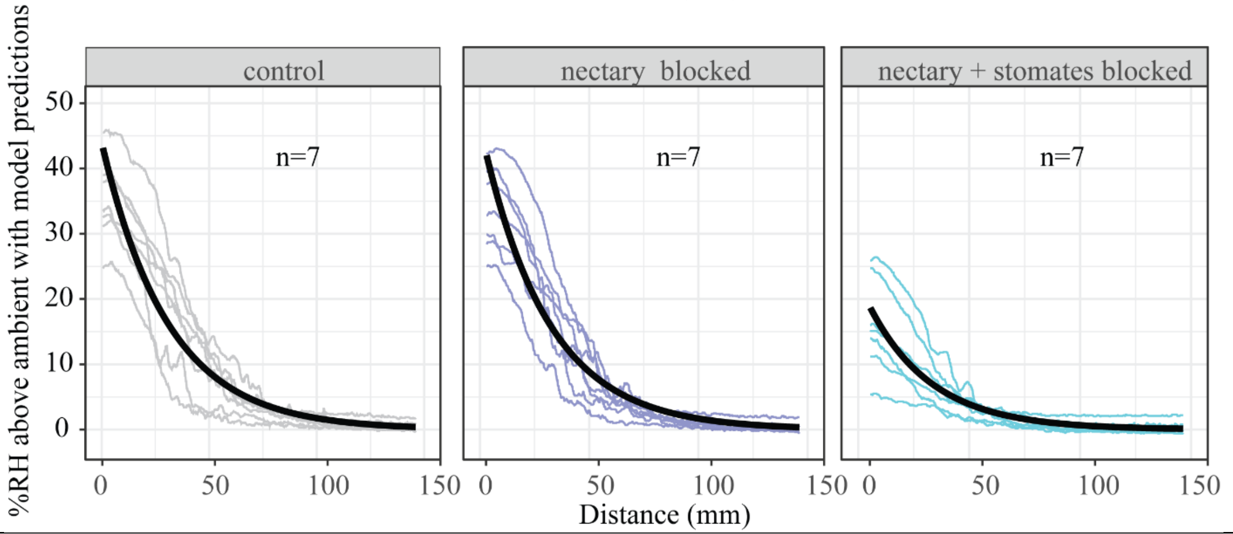


**Figure S4.** Model predictions (solid black line) for floral humidity of control and manipulated flowers. Individual flower transects (color-coded) are shown for the indicated treatment. Sample sizes are mentioned in each panel.

**Table S6.** Multiple comparisons of the model estimates for the **decay rate (α)** across the different treatments are shown in **figure S4** (Post hoc Tukey HSD).

| comparisons | estimate | SE | df | T ratio | P-value |
|---|---|---|---|---|---|
| control - nectary blocked | -0.00039462 | 0.00373999 | 13624 | -0.106 | 0.994 |
| control - (nectary+stomates blocked) | -0.00195809 | 0.00376801 | 13624 | -0.520 | 0.862 |
| nectary blocked - (nectary+stomates blocked) | -0.00156347 | 0.00376792 | 13624 | -0.415 | 0.909 |

**Table S7.** Multiple comparisons of the model estimates for the **intercept y0** across the different treatments shown in **figure S4** (Post hoc Tukey HSD).

| comparisons | estimate | SE | df | T ratio | P-value |
|---|---|---|---|---|---|
| control - nectary blocked | 1.14949123 | 4.40171314 | 13624 | 0.261 | 0.96310419 |
| control - (nectary+stomates blocked) | 24.6623467 | 4.40178785 | 13624 | 5.603 | 0.00000007 |
| nectary blocked - (nectary+stomates blocked) | 23.5128555 | 4.40180713 | 13624 | 5.342 | 0.00000028 |

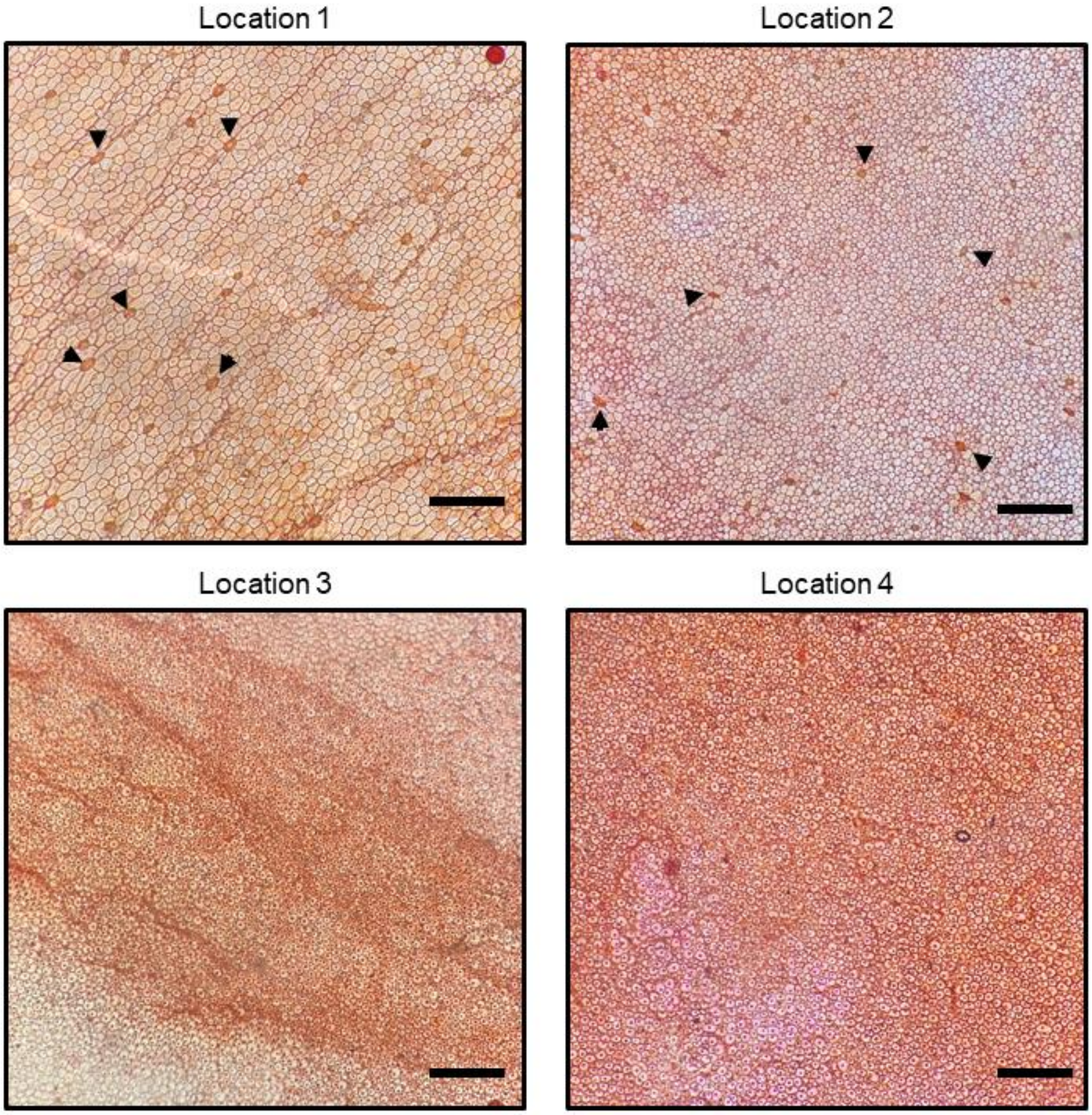


**Figure S5.** Pictures of the microscope slide mounted with safranin-stained flower peels from the inner surface of the flower (locations depicted in Figure 1E). Arrowheads point towards some of the stomates. No stomates were visible on Locations 3 and 4. Scale bar: 200 µm.

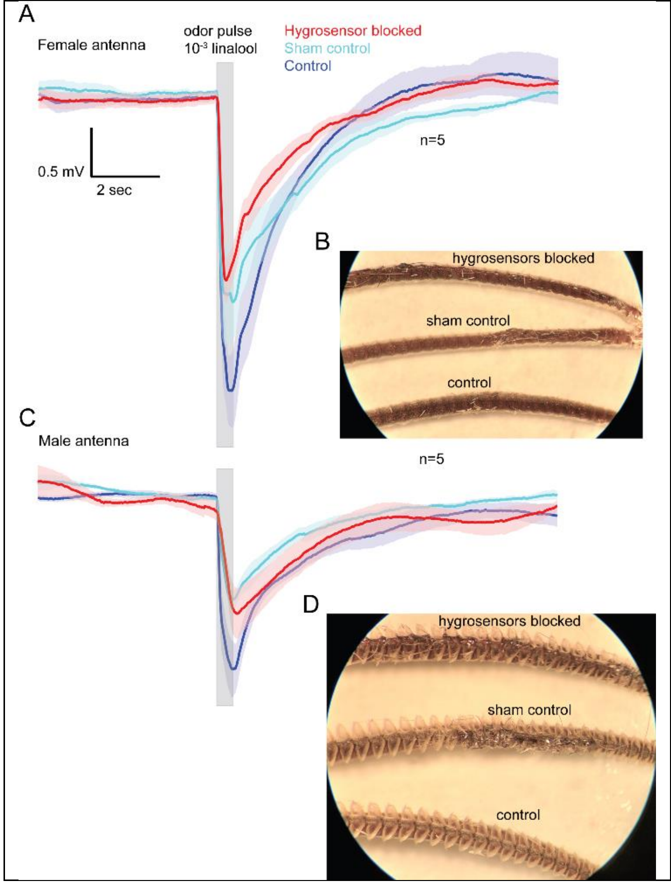


**Figure S6.** EAG responses of *M. sexta* for the indicated treatments to a 500ms odor pulse of $10^{-3}$ linalool.

A. Color coded mean (solid lines) and SEM (shaded area) responses of 5 female antenna for each treatment. Scale bar shows the amplitude and the duration of the response.
B. Representative images of the female antenna for each treatment.
C. Color coded mean (solid lines) and SEM (shaded area) responses of 5 male antenna for each treatment. Scale bar as above.
D. Representative images of the male antenna for each treatment

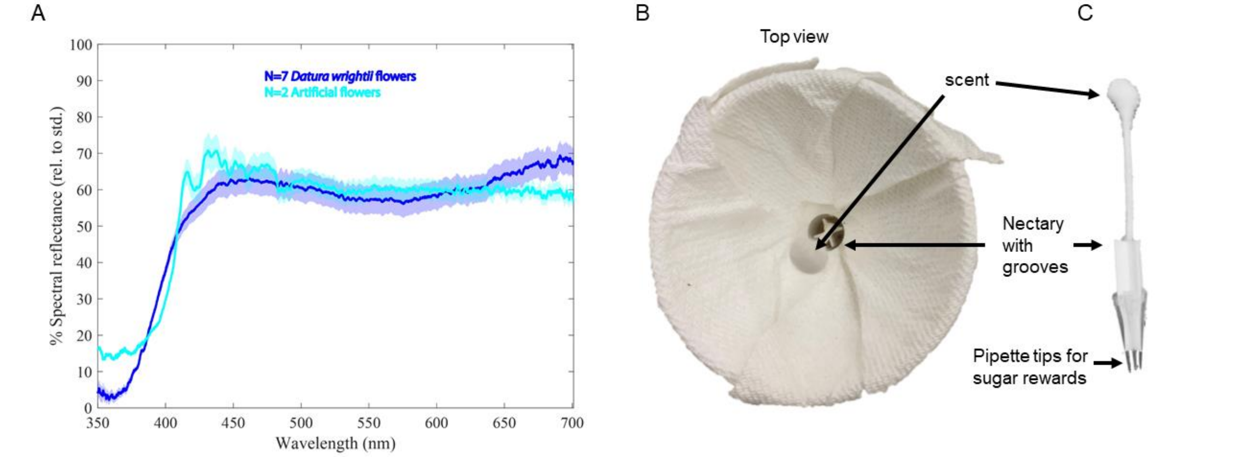


**Figure S7**. Design of the artificial flower used in the binary choice experiments.

A. Spectral reflectance of n=7 *Datura wrightii* flowers and that of the artificial flower n=2 (B) match closely with each other. Solid lines show the mean and shaded area shows the S.E.M
B. Top view of the artificial flower showing the four nectary grooves and the Q-tip occupying the space of style and stamens.
C. The internal assembly of the flower tube with the grooves ending in 4 pipette tips holding up to 50µl sugar solution each. 5µl Bergamot oil was added to the Q-tips at the start of each night of the experiment.

**A**

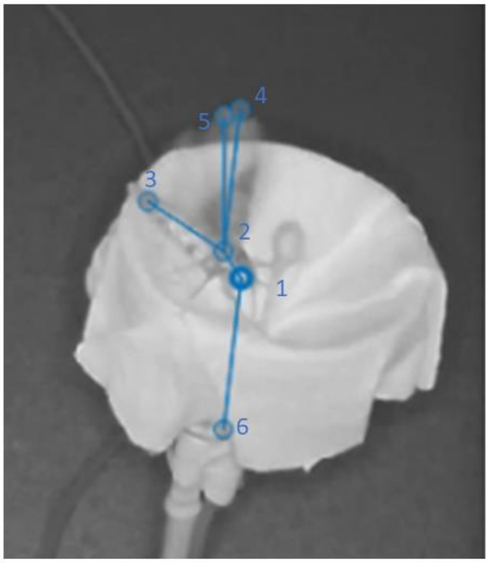


| Labeled Nodes |
|---|
| **1. Eye** |
| **2. Thorax-abdomen bridge** |
| **3. Abdomen tip** |
| **4. Left wing** |
| **5. Right wing** |
| **6. Proboscis tip** |

**B**

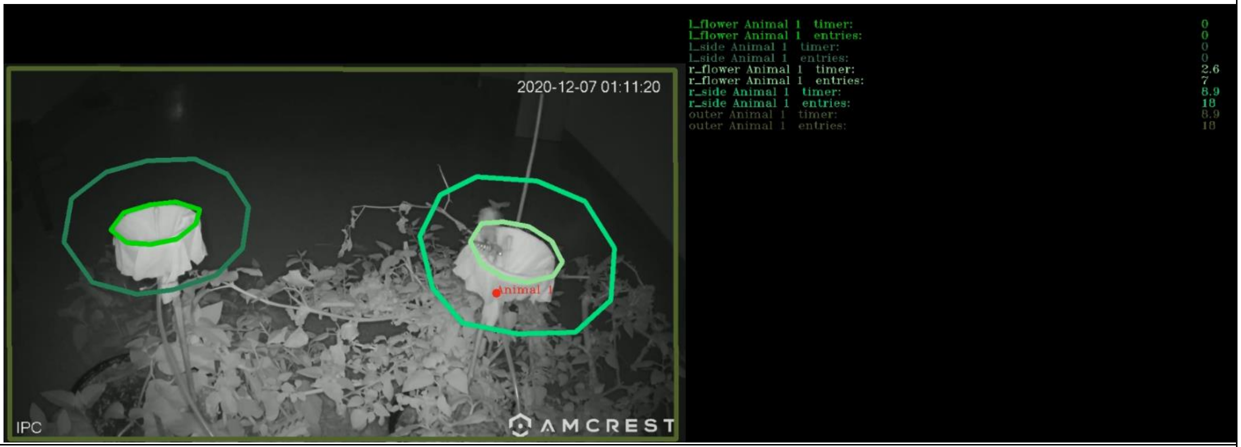


**Figure S8.** Methods for Behavior video analysis

(A) Moth body parts labeled using SLEAP v1.1.5, a deep-learning-based framework for estimating animal body part positions [1,2]

(B) Behavioral analysis of the SLEAP labeled moth body parts by tracking their frequency and time spent in the drawn region of interest (green circles) around flowers using SimBA, an open-source toolkit for behavioral analysis [3]
